# Humoral immunity to SARS-CoV-2 elicited by combination COVID-19 vaccination regimens

**DOI:** 10.1101/2022.05.13.491823

**Authors:** Zijun Wang, Frauke Muecksch, Friederike Muenn, Alice Cho, Shuai Zong, Raphael Raspe, Victor Ramos, Brianna Johnson, Tarek Ben Tanfous, Justin DaSilva, Eva Bednarski, Camila Guzman-Cardozo, Martina Turroja, Katrina G. Millard, Pinkus Tober-Lau, David Hillus, Kai-Hui Yao, Irina Shimeliovich, Juan Dizon, Anna Kaczynska, Mila Jankovic, Anna Gazumyan, Thiago Y. Oliveira, Marina Caskey, Paul D. Bieniasz, Theodora Hatziioannou, Florian Kurth, Leif Erik Sander, Michel C. Nussenzweig, Christian Gaebler

## Abstract

The SARS-CoV-2 pandemic prompted a global vaccination effort and the development of numerous COVID-19 vaccines at an unprecedented scale and pace. As a result, current COVID- 19 vaccination regimens comprise diverse vaccine modalities, immunogen combinations and dosing intervals. Here, we compare vaccine-specific antibody and memory B cell responses following two-dose mRNA, single-dose Ad26.COV2.S and two-dose ChAdOx1 or combination ChAdOx1/mRNA vaccination. Plasma neutralizing activity as well as the magnitude, clonal composition and antibody maturation of the RBD-specific memory B cell compartment showed substantial differences between the vaccination regimens. While individual monoclonal antibodies derived from memory B cells exhibited similar binding affinities and neutralizing potency against Wuhan-Hu-1 SARS-CoV-2, there were significant differences in epitope specificity and neutralizing breadth against viral variants of concern. Although the ChAdOx1 vaccine was inferior to mRNA and Ad26.COV2.S in several respects, biochemical and structural analyses revealed enrichment in a subgroup of memory B cell neutralizing antibodies with distinct RBD-binding properties resulting in remarkable potency and breadth.

## Introduction

Coronavirus disease-2019 (COVID-19) vaccine programs are a historic public health success that saved countless lives and prevented millions of Severe Acute Respiratory Syndrome Coronavirus (SARS-CoV-2) infections (Vilches et al., 2022). Vaccination is a multifaceted global effort involving a diverse collection of vaccine platforms including mRNA, adenoviral vector-based, inactivated virus and recombinant protein immunogens (Mathieu et al., 2021). Detailed evaluation of the different vaccine-specific immune responses will inform improved vaccination strategies for the prevention of COVID-19 and other respiratory viral infections of pandemic potential (Zhang et al., 2022).

With close to 2.5 billion administered doses, the ChAdOx1 nCoV-19 (AZD1222) vaccine accounted for over one third of all global COVID-19 vaccine doses administered in 2021 (Mallapaty et al., 2021; Mathieu et al., 2021). The ChAdOx1 vaccine encodes full-length wild-type (Wuhan-Hu-1) SARS-CoV-2 spike protein without the prefusion-stabilizing mutations found in the three US-approved vaccines (BNT162b2, mRNA-1273, and Ad26.COV2.S) (Watanabe et al., 2021). Outside of the US, ChAdOx1 received regulatory approval as a two-dose vaccine administered at an interval of 4–12 weeks. Unfortunately, ChAdOx1 vaccination was associated with immune thrombocytopenia, a rare but serious complication that has been described after administration of adenoviral vector vaccines. As a result, many individuals receiving a ChAdOx1 prime were subsequently boosted with an mRNA vaccine (Klok et al., 2022).

The combination (ChAdOx1/mRNA vaccine) prime-boost regimen showed enhanced immunogenicity (Barros-Martins et al., 2021; Hillus et al., 2021; Normark et al., 2021; Schmidt et al., 2021b), however both ChadOx1-based vaccine regimens proved to be effective with substantial protection against COVID-19 hospitalization and death (Andrews et al., 2022; Nordstrom et al., 2021).

In-depth analyses of antibody and memory B cell responses after natural infection, mRNA (BNT162b2, mRNA-1273) and Janssen Ad26.COV2.S vaccination have been performed (Cho et al., 2021; Gaebler et al., 2021; Muecksch et al., 2022; Robbiani et al., 2020; Wang et al., 2021c; Wang et al., 2021d). However, far less is known about the responses elicited by the ChadOx1 vaccine even though it was used in more countries that any other COVID-19 vaccine (Mathieu et al., 2021). Here, we compare vaccine-specific antibody and memory B cell responses to 2-dose mRNA (BNT162b2 or mRNA-1273), 1-dose Janssen Ad26.COV2.S, 2-dose ChAdOx1 (AZ/AZ) or ChAdOx1/BNT162b2 combination (AZ/BNT) vaccines.

## Results

Four cohorts of study participants with different vaccination regimens were recruited and sampled prospectively. All cohorts included sampling time points at approximately 1 and 6 months after the 1^st^ vaccine dose. An additional sampling time point at 1 month after 2^nd^ vaccination was available for the mRNA (1.3m after 2^nd^ dose=2.3m after 1^st^ dose), AZ/BNT and AZ/AZ (1m after 2^nd^ dose=4m after 1^st^ dose) 2-dose vaccination regimens. The vaccination and blood collection schedule for all cohorts in this study is depicted in Fig. 1a.

**Fig. 1:**
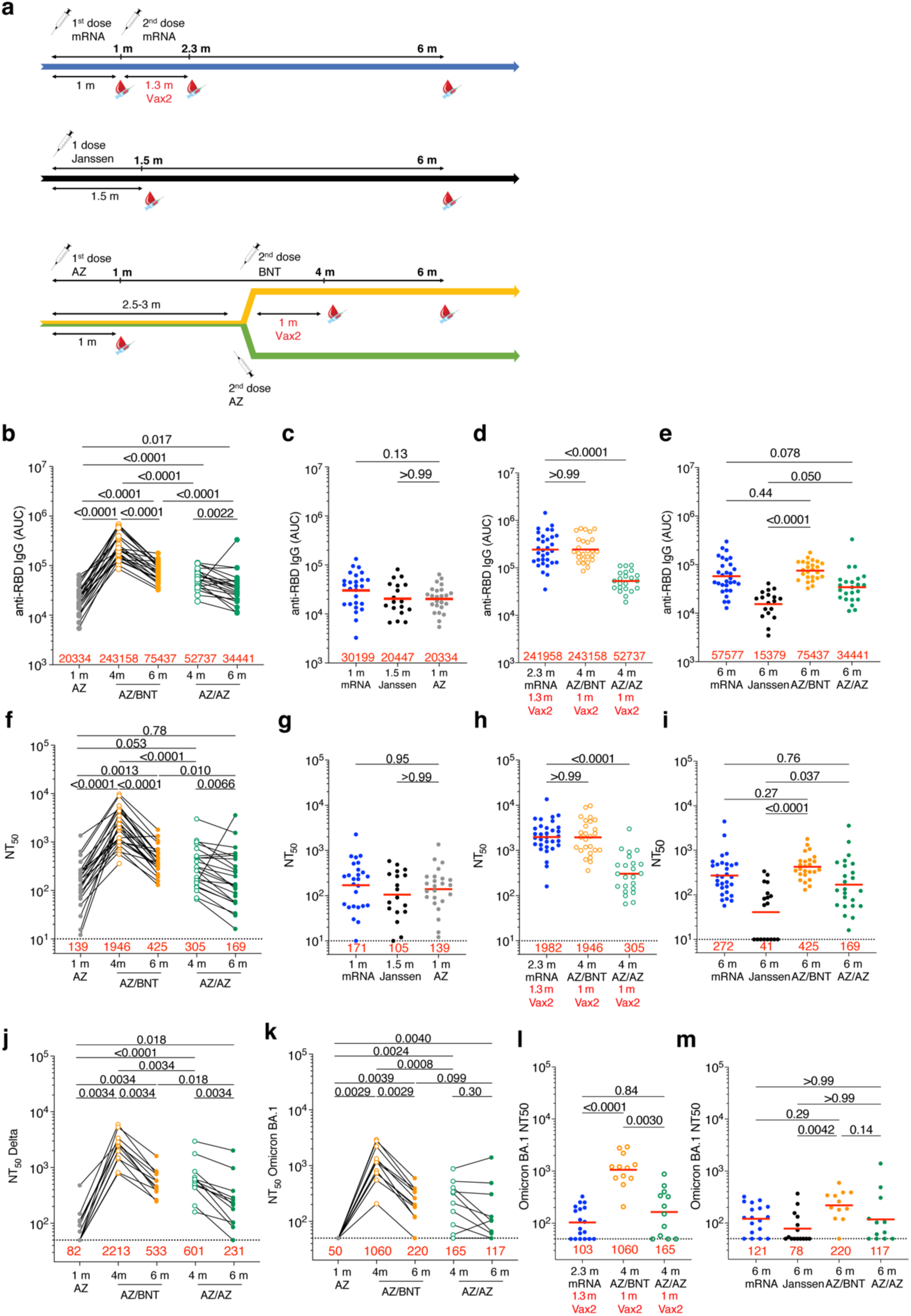
Plasma binding and neutralizing activity. **a,** Vaccination and blood donation schedules for mRNA vaccinees (upper panel), Ad26.COV.2S (Janssen) vaccinees (middle panel), and ChAdOx1 (AZ) vaccinees boosted with either BNT162b2 (BNT, upper half of lower panel) or AZ (lower half of lower panel). **b,** Area under the curve (AUC) for plasma IgG antibody binding to SARS-CoV-2 Wuhan-Hu-1 RBD 1 month (m) after mRNA prime (Cho et al., 2021), or Janssen Ad26.COV.2S prime (Cho et al., 2022) or AZ prime, as well as 4 months or 6 months after the initial AZ prime (AZ/BNT; n=26) or (AZ/AZ; n=23). Lines connect longitudinal samples. **c-e,** AUC for plasma IgG binding to Wuhan-Hu-1 RBD in vaccinees 1 m after AZ prime compared to mRNA prime (Cho et al., 2021) or Janssen Ad26.COV2.S (Cho et al., 2022) prime at similar timepoint **(c)**, mRNA vaccinees 2.3 m after initial dose (Cho et al., 2021) compared to AZ/BNT and AZ/AZ vaccinees 4 m after initial dose **(d)**, or mRNA vaccinees 6 m after initial dose (Cho et al., 2021) and Janssen Ad26.COV2.S vaccinees 6 m after one dose (Cho et al., 2022) compared to AZ/BNT and AZ/AZ vaccinees 6 m after initial dose **(e)**. **f-i,** Anti-SARS-CoV-2 NT_50_s of plasma measured by a SARS-CoV-2 pseudotype virus neutralization assay using wild-type (Wuhan-Hu-1 (Wu et al., 2020)) SARS-CoV-2 pseudovirus (Robbiani et al., 2020; Schmidt et al., 2020) in plasma samples shown in panels **a-e**. **j-m,** Plasma neutralizing activity against indicated SARS-CoV-2 Delta **(j)** and Omicron **(k)** variants for n=24 (AZ/BNT: n=12 and AZ/AZ:n=12) randomly selected samples assayed in HT1080Ace2 cl.14 cells. **l-m,** mRNA vaccinees 2.3 m after initial dose (Cho et al., 2021) compared to AZ/BNT and AZ/AZ vaccinees 4 m after initial dose **(l)**, or mRNA vaccinees 6 m (Cho et al., 2021) and Janssen Ad26.COV2.S vaccinees 6 m after initial dose (Cho et al., 2022) compared to AZ/BNT and AZ/AZ vaccinees 6 m after initial dose **(m)**. See Methods for a list of all substitutions/deletions/insertions in the spike variants. Deletions/substitutions corresponding to viral variants were incorporated into a spike protein that also includes the R683G substitution, which disrupts the furin cleavage site and increases particle infectivity. All experiments were performed at least in duplicate. Red bars and values represent geometric mean values. Statistical significance was determined by two-tailed Mann-Whitney test for unpaired observations or by Wilcoxon matched-pairs signed rank test for paired observations followed by Holm-Šídák test for multiple comparisons (**a**, **f, j-k**), two-tailed Kruskal-Wallis test with subsequent Dunn’s multiple comparisons (**b-e, g-I, l-m**).

For the AZ/BNT and AZ/AZ cohort a total of 49 health-care workers with no prior history of SARS-CoV-2-infection who received a ChAdOx1 vaccine prime followed by ChAdOx1 or BNT162b2 boost 10–12-weeks later were enrolled in a prospective observational cohort study in Berlin (Germany) (Hillus et al., 2021). 23 and 26 study participants received ChAdOx1 or mRNA as a second dose, respectively. Volunteers ranged in age from 20-65 years and were 65% female. For detailed demographic information, see Methods and Table S1 and (Cho et al., 2022; Muecksch et al., 2022).

### Plasma binding and Neutralization

Plasma antibody binding titers to SARS-CoV-2 RBD were measured by enzyme-linked immunosorbent assays (ELISA) (Cho et al., 2021; Wang et al., 2021c). RBD-binding IgG levels 1 month after ChAdOx1 prime were lower but not significantly different to antibody levels following a single dose of an mRNA vaccine (Cho et al., 2021) or the Janssen Ad26.COV2.S vaccine (Cho et al., 2022) at similar time points (Fig. 1b-c). ChAdOx1 and mRNA boosting enhanced IgG titers 12-fold (AZ/BNT) and 2.6-fold (AZ/AZ) 1 month after the 2^nd^ vaccine dose, respectively (p<0.0001, Fig. 1b, d). In both cases, anti-RBD antibodies in plasma decreased significantly between 4 and 6 months (AZ/BNT: 3.2-fold, p<0.0001; AZ/AZ: 1.5-fold, p=0.0022, Fig.1b), but antibodies binding to RBD following combination AZ/BNT vaccination remained significantly higher 6 months after the initial priming dose (p<0.0001, Fig. 1b). Anti-RBD IgG levels after the AZ/BNT boost were directly correlated with initial antibody levels after the prime (Fig. S1a, r=0.50, p=0.012). Consistent with other reports (Barros-Martins et al., 2021; Kaku et al., 2022; Pozzetto et al., 2021), AZ/BNT vaccinees exhibited anti-RBD plasma reactivity 6 months after the initial prime that were comparable to individuals who received two doses of an mRNA vaccine (Fig. 1d, e). By contrast, antibody levels following AZ/AZ vaccination remained substantially lower compared to individuals who received two doses of an mRNA vaccine.

Nevertheless, individuals who received AZ/AZ vaccination showed serum antibody levels that exceeded those of Janssen Ad26.COV2.S vaccinees at 6m post-vaccination (Fig. 1e). In contrast to IgG, AZ/BNT and AZ/AZ vaccination induced similar IgM and IgA anti-RBD antibody levels (Fig. S1b-c).

RBD-binding IgG titers were negatively correlated with age 4 month after the initial prime for AZ/AZ, but not AZ/BNT vaccination (r=-0.51, *P*=0.015, Fig. S1d-e). There were no sex-related differences in antibody levels following AZ/BNT or AZ/AZ vaccination (Fig. S1f). Notably, 4 months after the initial prime, antibody levels in AZ/BNT vaccinees were negatively correlated with the interval between prime and 2^nd^ vaccination, suggesting that earlier administration of a heterologous booster vaccination may result in optimal protection (r=-0.50, *P*=0.010, Fig. S1g- h).

Neutralizing activity was determined for the same participants, using HIV-1 pseudotyped with Wuhan-Hu-1 SARS-CoV-2 spike (S) protein (Robbiani et al., 2020; Schmidt et al., 2020) (Table S1).

The geometric mean half-maximal neutralizing titer (NT50) 1 month after the ChAdOx1 initial prime were comparable to a single dose of an mRNA vaccine (Cho et al., 2021) or Janssen Ad26.COV2.S (Cho et al., 2022) vaccine (Fig. 1f, g). Administration of a second dose increased NT50s among AZ/BNT and AZ/AZ vaccinees from 139 to 1946 and 305, respectively (p<0.0001, Fig.1f). In line with the greater initial neutralizing activity, the decrease between 4-6 months after the initial prime was more pronounced among combination AZ/BNT than AZ/AZ vaccinees (4.6-fold, p<0.0001 vs. 1.8- fold, p=0.0066 respectively, Fig.1f). Nevertheless, compared to AZ/AZ vaccinees, plasma neutralizing activity remained significantly higher 6 months after the initial prime in AZ/BNT vaccinees (p=0.01, Fig. 1f)

Consistent with ELISA reactivity, AZ/BNT elicited similar neutralizing activity as two doses of an mRNA vaccine 6 months after the initial prime (Fig. 1h). By contrast, plasma neutralizing activity after AZ/AZ vaccination was substantially lower than in individuals who received two doses of an mRNA vaccine but exceeded neutralizing titers of individuals that received a single dose of the Janssen Ad26.COV2.S vaccine (Fig. 1h, i).

Plasma neutralizing activity for 24 randomly selected samples (n=12, AZ/BNT; n=12, AZ/AZ) was also assessed against SARS-CoV-2 Delta and Omicron BA.1 variants using viruses pseudotyped with appropriate variant spikes (Cho et al., 2021; Schmidt et al., 2021a; Schmidt et al., 2020; Wang et al., 2021d). Four months after the initial AZ/BNT prime vaccination, neutralizing titers against Delta and Omicron BA.1 were 5.5-, and 11.6-fold lower than against Wuhan-Hu-1, with a further decrease to 5.6- and 13.6-fold lower activity at the 6-month time point respectively (Fig. 1j-k and Fig. S1i). Similarly, AZ/AZ vaccination resulted in 5.8- and 21- fold lower neutralizing activity against Delta and Omicron BA.1 than against Wuhan-Hu-1 respectively at the 4-month time point. While Delta neutralization further decreased 6.4-fold compared to Wuhan-Hu-1 at the 6-month time point, the neutralizing activity against Omicron BA.1, which was initially very low, decreased to a lesser extent among AZ/AZ vaccinees (Fig. 1j-k, Fig. S1i-k).

Remarkably, 1 month after the 2^nd^ vaccine dose Omicron BA.1 neutralizing titers in combination AZ/BNT vaccinees exceeded neutralizing activity after AZ/AZ or 2-dose mRNA vaccination at similar time points by 6.4- and 10.3-fold, respectively (p=0.003 and p<0.0001 Fig. 1l). Omicron BA.1 neutralizing titers remained higher in AZ/BNT vaccinees, but were not statistically different from mRNA, or AZ/AZ vaccinees 6 months after the prime, while titers in Janssen Ad26.COV2.S vaccinees were significantly lower (Fig. 1m).

### Memory B cell responses to SARS-CoV-2 RBD and NTD

To compare the development of B cell memory after COVID-19 vaccination, we initially enumerated memory B cells expressing surface receptors binding to the Receptor-Binding Domain (RBD) and N-Terminal Domain (NTD) of the SARS-CoV-2 spike protein using fluorescently labeled proteins (Fig. 2a, Fig. S2a-d). The number of RBD-binding memory B cells found in circulation 1 month after AZ prime was significantly lower than after mRNA (p<0.0001, (Bednarski et al., 2022; Cho et al., 2021)) and Janssen Ad26.COV2.S vaccination (p=0.0029 (Cho et al., 2022)) (Fig. 2b). Although the number of RBD-binding memory B cells increased after AZ/BNT or AZ/AZ boosting (Fig. 2c), the number remained lower than after 2- dose mRNA vaccination (AZ/BNT: p=0.02, AZ/AZ: p=0.0003, Fig. 2d). By contrast, the number of NTD-binding memory B cells remained unchanged after the 2^nd^ dose (Fig. S2c-d) and was similar to 2-dose mRNA and significantly lower than in Janssen Ad26.COV2.S vaccinees 6 months post vaccination (AZ/BNT: p=0.0007; AZ/AZ: p=0.0001, Fig. S2e).

**Fig. 2:**
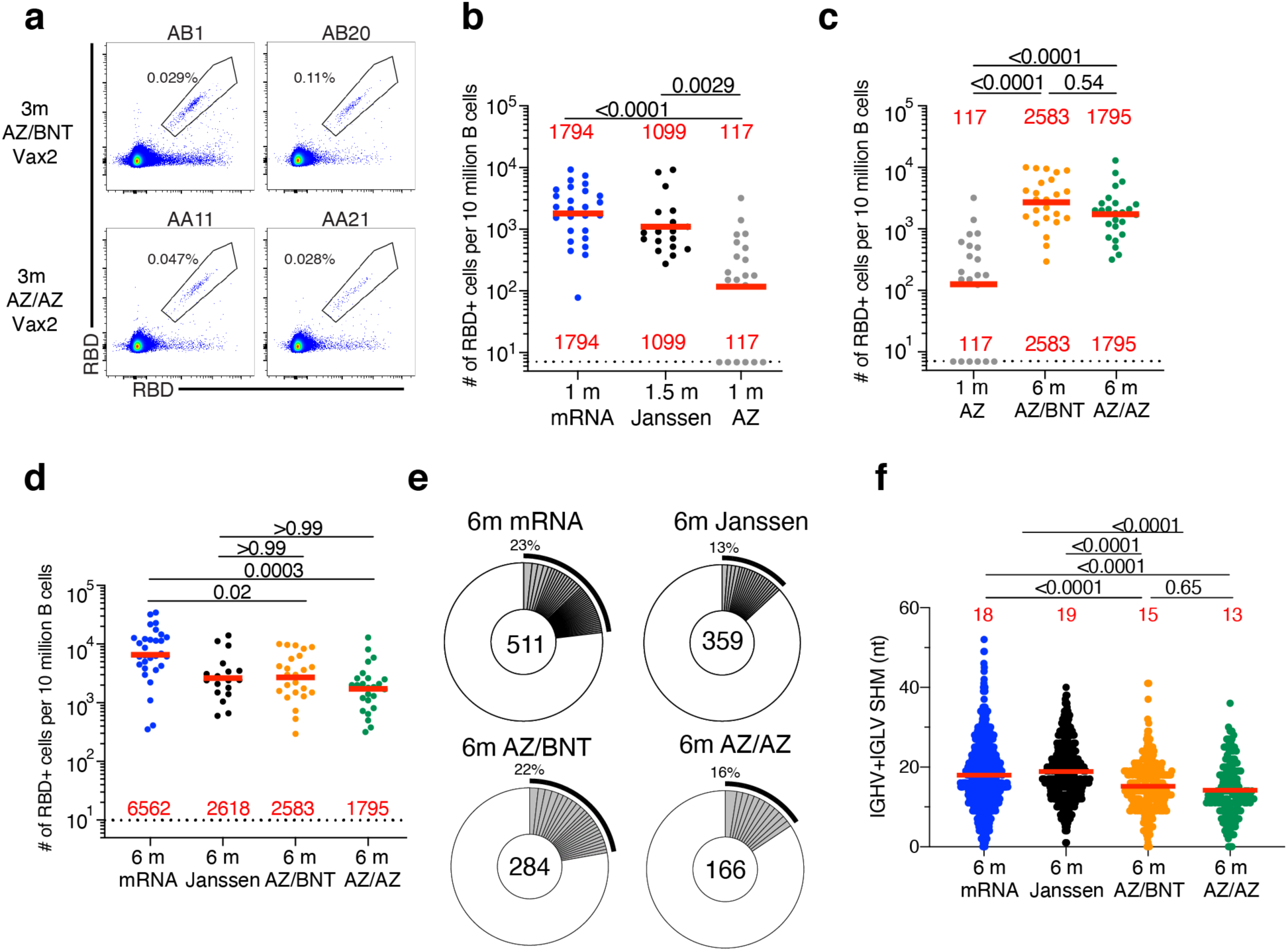
Anti-SARS-CoV-2 RBD B cell memory. **a,** Representative flow cytometry plots showing dual AlexaFluor-647- and PE-Wuhan-Hu-1-RBD-binding single sorted B cells from ChAdOx1/BNT162b2 (AZ/BNT, n=2) and ChAdOx1/ChAdOx1 (AZ/AZ, n=2) vaccinees 6 months (m) after initial dose. Gating strategy shown in **Fig. S2**. Percentage of antigen-specific B cells is indicated. **b,** Number of Wuhan-Hu-1 RBD-specific B cells per 10 million (M) B cells in mRNA vaccinees 1 m after prime (Cho et al., 2021) and Ad26.COV.2S (Janssen) vaccinees 1.5 m after prime (Cho et al., 2022) compared to AZ vaccinees 1 m after prime. **c,** Number of Wuhan- Hu-1 RBD-specific B cells per 10 M B cells for AZ vaccinees 1 m after prime compared to AZ/BNT and AZ/AZ vaccinees 6 m after initial dose. **d,** Number of Wuhan-Hu-1 RBD-specific B cells per 10 M B cells for mRNA vaccinees 6 m after initial dose (Cho et al., 2021) and Janssen Ad26.COV2.S vaccinees 6 m after prime (Cho et al., 2022) compared to AZ/BNT and AZ/AZ vaccinees 6 m after initial dose. **e,** Pie charts show the distribution of antibody sequences obtained from Wuhan-Hu-1 RBD-specific memory B cells of mRNA vaccinees 6 m after initial dose and Ad26.COV.2S (Janssen) vaccinees 6 m after initial prime, or AZ/BNT and AZ/AZ vaccinees 6 m after initial dose. The number inside the circle indicates the aggregate number of sequences analyzed for each cohort. Grey slices indicate expanded clones (same *IGHV* and *IGLV* genes with highly similar CDR3s, see Methods) found within the same individual. Pie slice size is proportional to the number of clonally related sequences. The black outline and associated numbers indicate the total percentage of clonally expanded sequences. **f,** Number of nucleotide somatic hypermutations (SHM) in *IGHV* + *IGLV* sequences obtained from Wuhan-Hu-1 RBD- specific memory B cells of mRNA vaccinees 6 m after initial dose and Janssen Ad26.COV2.S vaccinees 6 m after prime compared to AZ/BNT and AZ/AZ vaccinees 6 m after initial dose. Red bars and numbers in **b, c**, and d represent geometric mean value, and in f represent mean values. Statistical difference in **b, c, d** and **f**, was determined by two-tailed Kruskal Wallis test with subsequent Dunn’s multiple comparisons.

To examine the specificity and neutralizing activity of the antibodies produced by memory cells we purified single Wuhan-Hu-1 RBD-specific B cells, sequenced their antibody genes, and produced the recombinant antibodies *in vitro*. 450 paired anti-RBD antibody sequences were obtained from 22 vaccinees (AZ/BNT n=10; AZ/AZ n=12) sampled 6 months after the initial prime (Fig. 2e, and Fig. S2f-g, Fig. S3, Table S2). Clonally expanded RBD-specific B cells across the different vaccine regimens 6 months after prime represented 23%, 13%, 22% and 16% of all memory cells from mRNA, Janssen Ad26.COV2.S, AZ/BNT and AZ/AZ vaccinees, respectively (Fig. 2e). Similar to mRNA and Janssen Ad26.COV2.S vaccinees, VH3-30, VH1-46 and VH3-53 genes were overrepresented among AZ/BNT and AZ/AZ vaccinees (Fig. S4, (Cho et al., 2021; Cho et al., 2022)). There was no difference in the number of somatic mutations between AZ/BNT or AZ/AZ vaccinees, although both groups showed significantly lower accumulation of somatic mutations than mRNA or Janssen Ad26.COV2.S vaccinees assayed at similar timepoints (p<0.0001, Fig. 2f).

We conclude that there are substantial differences in the magnitude, clonal composition, and antibody maturation of the RBD-specific memory B cell compartment between the different vaccination regimens. However, homologous and combination booster vaccination after a ChAdOx1 prime are not significantly different with respect to the number of memory B cells that develop, or their levels of somatic mutation.

### Neutralizing activity of monoclonal antibodies

Next, we compared the neutralizing activity of monoclonal antibodies elicited by mRNA, Janssen Ad26.COV2.S, and AZ/BNT or AZ/AZ vaccination. 291 anti-RBD monoclonal antibodies were expressed and tested for binding by ELISA. 94% (n=277) bound to the Wuhan-Hu-1 RBD, indicating the high efficiency of RBD-specific memory B cell isolation (Table S3). The geometric mean ELISA half-maximal concentration (EC_50_) of the RBD-binding monoclonal antibodies elicited by AZ/AZ or AZ/BNT vaccination was comparable (Fig. 3a). EC_50_s represent an indirect measure of affinity. To directly examine anti-RBD antibody affinity, we performed biolayer interferometry (BLI) experiments on a subset of antibodies (n=66 from AZ/BNT and n=62 from AZ/AZ). Affinity was comparable among antibodies elicited by mRNA, Janssen Ad26.COV2.S, and AZ/BNT or AZ/AZ vaccination (Fig. 3b, (Cho et al., 2021)). All 277 RBD-binding IgG monoclonal antibodies were tested for neutralization against viruses pseudotyped with Wuhan-Hu-1 SARS-CoV-2 spike protein (183 and 94 antibodies isolated from AZ/BNT and AZ/AZ vaccinees, respectively). Memory B cell antibodies elicited by mRNA, Janssen Ad26.COV2.S, AZ/BNT and AZ/AZ vaccination 6 months after the prime showed comparable activity (Fig 3c). Similarly, the proportion of neutralizing to non-neutralizing antibodies for all four regimens was not significantly different (Fig. 3d). We conclude that memory B cells present in circulation 6 months after initial mRNA, Janssen Ad26.COV2.S, AZ/AZ and AZ/BNT vaccine doses express antibodies with similar binding affinities and neutralizing potency against Wuhan-Hu-1 SARS-CoV-2.

**Fig. 3:**
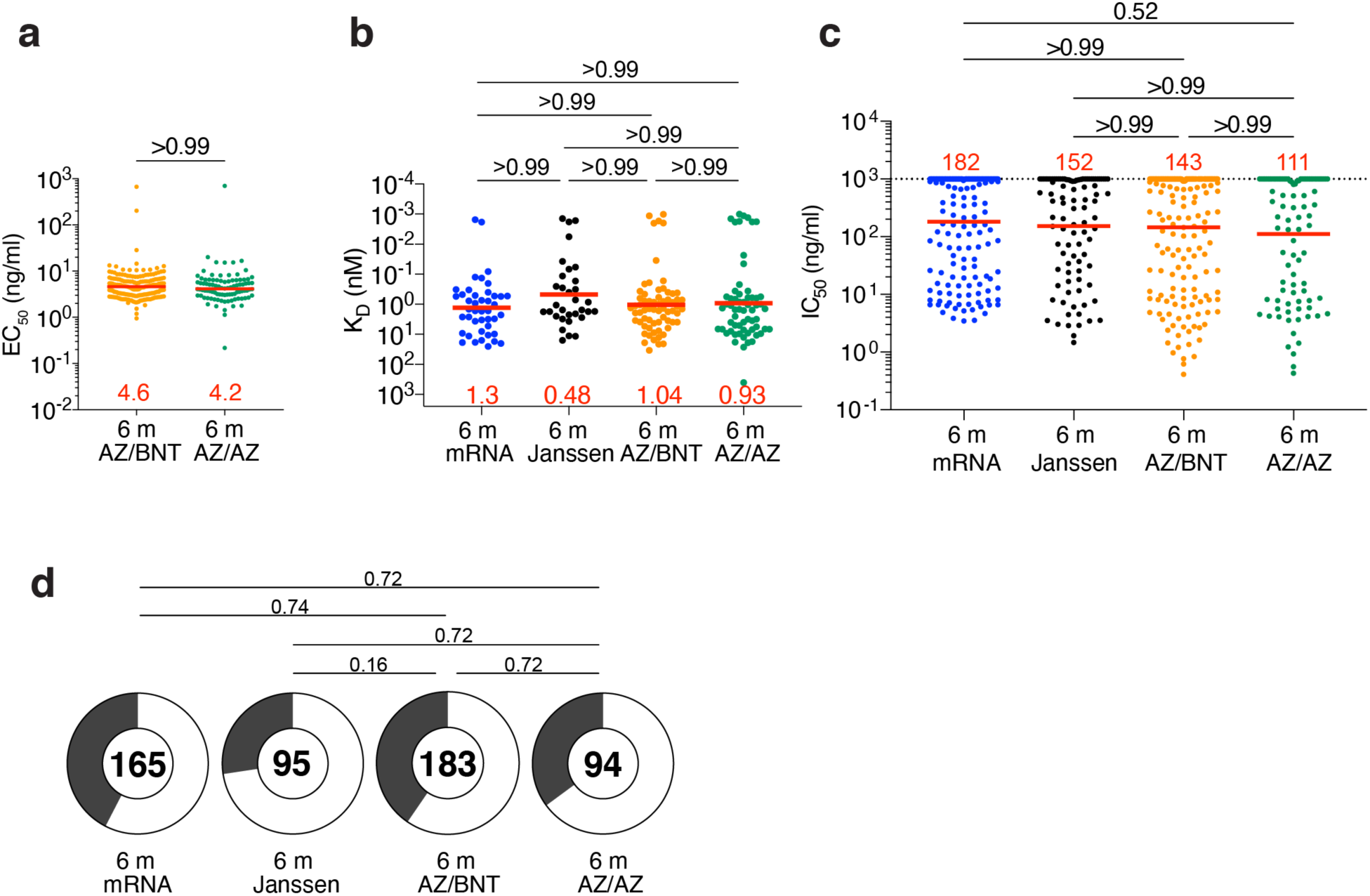
Anti-SARS-CoV-2 monoclonal antibodies. **a**, Graph shows half-maximal effective concentration (EC_50_) of n=277 Wuhan-Hu-1 RBD-binding monoclonal antibodies (mAbs) measured by ELISA against Wuhan-Hu-1 RBD from AZ/BNT (n=183) and AZ/AZ (n=94) vaccinees. **b**, Graph showing affinity measurements (K_D_s) for Wuhan- Hu-1 RBD measured by BLI for antibodies cloned from mRNA vaccinees 6 months(m) after initial dose (n=43) (Cho et al., 2021), from Janssen Ad26.COV2.S vaccinees 6 m (n=33) after prime (Cho et al., 2022), compared to antibodies cloned from AZ/BNT (n=189) and AZ/AZ (n=94) vaccinees 6 m after initial dose. **c**, Graphs show anti-SARS-CoV-2 neutralizing activity of monoclonal antibodies measured by a SARS-CoV-2 pseudotype virus neutralization assay using wild-type (Wuhan-Hu-1(Wu et al., 2020)) (SARS-CoV-2 pseudovirus (Robbiani et al., 2020; Schmidt et al., 2020)) for antibodies cloned from mRNA vaccinees and 6 m after initial dose (n=262) (Cho et al., 2021), or from Janssen Ad26.COV2.S vaccinees (n=95) 6 m after prime (Cho et al., 2022), compared to antibodies cloned from AZ/BNT (n=189) and AZ/AZ (n=94) vaccinees 6 m after initial dose. **d**. Pie charts indicated the frequency of neutralizing (IC_50_<1000 ng/mL, white) vs. non-neutralizing (IC_50_>1000 ng/mL, black) antibodies cloned from mRNA vaccinees(Cho et al., 2021), Janssen Ad26.COV2.S vaccinees(Cho et al., 2022), AZ/AZ vaccinees and AZ/BNT vaccinees. Red bars and lines indicate geometric mean values. Statistical significance in a, b, and c was determined by two-tailed Kruskal Wallis test with subsequent Dunn’s multiple comparisons. Pie charts were compared using a two-tailed Fisher’s exact test.

### Epitopes and Neutralizing Breadth

SARS-CoV-2 infection and vaccination elicit anti-RBD antibodies that target four structurally defined classes of epitopes on the RBD (Barnes et al., 2020; Muecksch et al., 2022; Muecksch et al., 2021; Wang et al., 2021c; Yuan et al., 2020). Class 1 and 2 antibodies block ACE2 binding directly, and Class 3 and 4 antibodies target more conserved regions on the RBD (Gaebler et al., 2021; Muecksch et al., 2022; Muecksch et al., 2021; Wang et al., 2021c). Class 1 and 2 antibodies develop early after infection or mRNA-immunization (Muecksch et al., 2022), while Janssen Ad26.COV2.S vaccination leads to a more diverse, early memory B cell response that is dominated by Class 3 and 4 antibodies (Cho et al., 2022). Nevertheless, continued memory B cell evolution results in comparable epitope specificities 5-6 months after the initial mRNA or Janssen Ad26.COV2.S immunization (Cho et al., 2022).

To define the epitopes recognized by anti-RBD memory antibodies elicited by AZ/BNT or AZ/AZ vaccination, we performed BLI competition experiments. A preformed antibody-RBD-complex was exposed to a second antibody targeting one of four classes of structurally defined epitopes (Barnes et al., 2020; Robbiani et al., 2020) (C105 as Class 1; C144 as Class 2; C135 as Class 3; and CR3022 as Class 4). We examined 128 RBD-binding antibodies randomly obtained from the AZ/BNT (n=66) and AZ/AZ (n=62) vaccinees. This included AZ/BNT (n=44) and AZ/AZ (n=39) antibodies with IC_50_s less than 1000 ng/mL.

The epitope distribution of the memory antibody repertoires was significantly different between all four vaccine regimens (Fig. S5a). Moreover, the overall epitope specificities of the antibody repertoires were significantly different between mRNA vaccinees and AZ/BNT or AZ/AZ vaccinees (Fig. 4a). This was particularly evident among neutralizing (IC_50_<1000ng/ml) antibodies for which the frequency of antibodies that target unknown epitopes (non-classified) was highly enriched in the antibody repertoire isolated from AZ/BNT or AZ/AZ vaccinees (Fig. 4a). At the same time, there were no significant differences in epitope specificities for non-neutralizing (IC_50_>1000 ng/ml) antibodies.

**Fig. 4:**
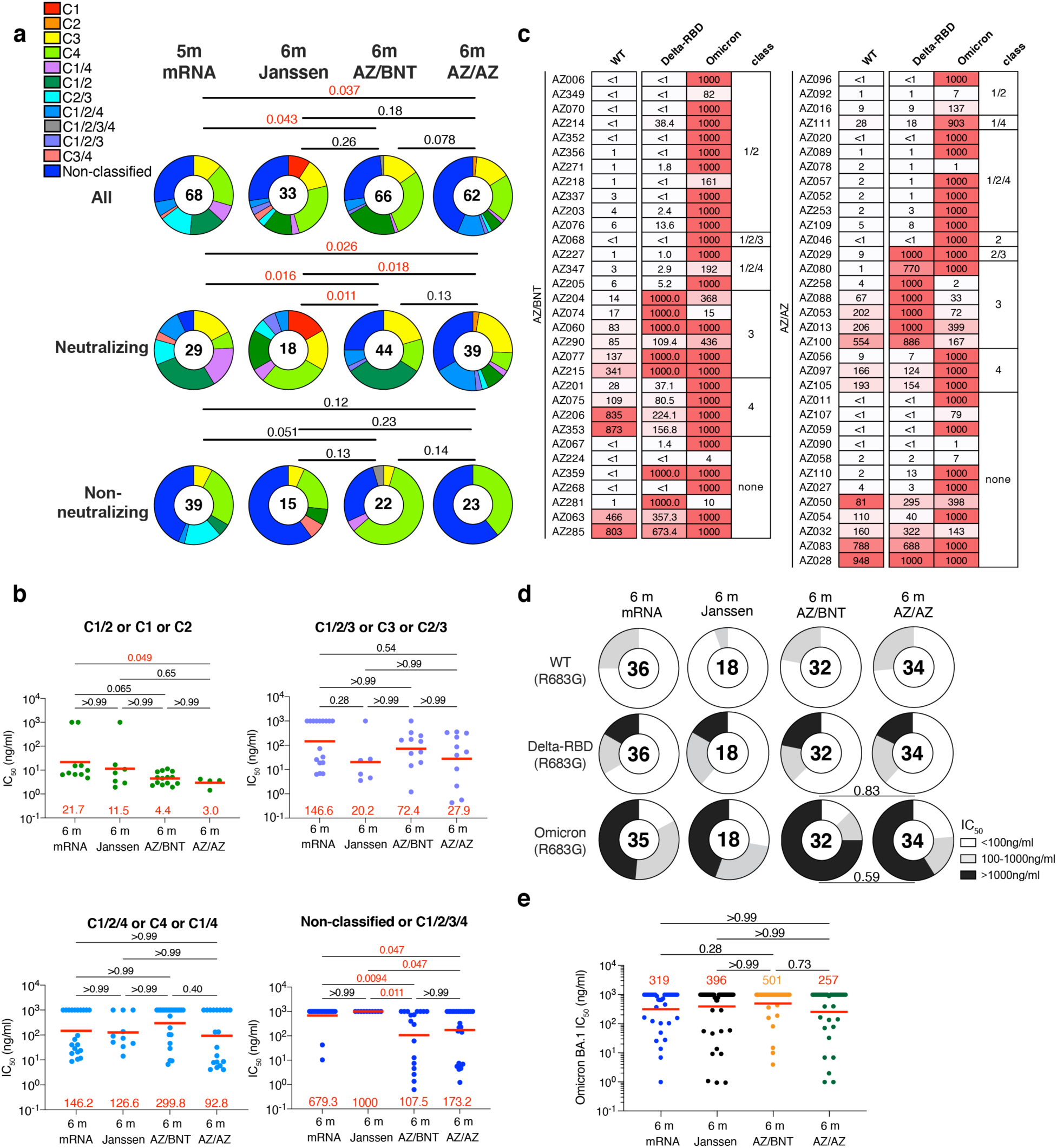
Epitopes and neutralizing breadth. Results of epitope mapping performed by competition BLI, comparing mAbs cloned from Janssen vaccinees 6 m (n=33) after prime (Cho et al., 2022) and mAbs cloned from mRNA vaccinees 6 m after initial dose (n=68) (Cho et al., 2021), to mAbs cloned from AZ/AZ (n=62) or AZ/BNT(n=66) vaccinees 6 m after initial dose. **a**, Pie charts show the distribution of the antibody classes among all RBD-binding antibodies, Wuhan-Hu-1 neutralizing antibodies only or non-neutralizing antibodies only. Statistical significance was determined by using a two-tailed Chi-square test. **b**, Graphs showing IC_50_ neutralization activity of antibodies indicated in **a** and **Fig. S5a**, with 4 categories by combining 1) C1/2 or C1 or C2 as C1/2; 2) C1/2/3 or C3 or C2/3 as C1/2/3; 3) C1/2/4 or C4 or C1/4 as C1/2/4; 4) Non-classified and C1/2/3/4 as non-classified. **c**, Heat-maps show IC_50_s of antibodies obtained from AZ/BNT vaccinees (n=32), and AZ/BNT vaccinees (n=34), against indicated mutant and variant SARS-CoV-2 pseudoviruses listed across the top. Delta-RBD indicate the L452R/T478K and Omicron BA.1. The deletions/substitutions corresponding to viral variants were incorporated into a spike protein that also includes the R683G substitution, which disrupts the furin cleavage site and increases particle infectivity. **d**. Pie charts show fraction of potent neutralizing (IC_50_<100ng/ml), less potent neutralizing (100ng/ml<IC_50_<1000ng/ml) and non-neutralizing (IC_50_>1000 ng/ml) antibodies in white, light and dark grey, respectively, for indicated SARS-CoV-2 pseudoviruses. Number in inner circles indicates number of antibodies tested. **e**. Graphs showing IC_50_ neutralization activity of antibodies mAbs cloned from Janssen Ad26.COV2.S vaccinees at 6 m (n=54) after prime (Cho et al., 2022) and mAbs cloned from mRNA vaccinees at 6 m after initial dose (n=35) (Cho et al., 2021), to mAbs cloned from AZ/AZ (n=34) or AZ/BNT(n=32) vaccinees 6 m after initial dose, against Omicron BA.1. Red bars and lines indicated geometric mean values. Statistical significance in **a** was determined by two-tailed Kruskal Wallis test with subsequent Dunn’s multiple comparisons in **b** and **e**. Statistical significance was determined using a two-tailed Chi-square test.

To examine the contribution of the different antibody classes to the neutralizing potency and breadth elicited by each of the four vaccine regimens, we regrouped the antibodies as follows: 1) Antibodies targeting Class 1 and/or 2 epitopes; 2) antibodies additionally or exclusively targeting Class 3 epitopes; 3) antibodies additionally or exclusively targeting Class 4 epitopes; or 4) non-classifiable antibodies. While neutralizing potency of the first 3 groups was comparable among all four vaccines regimens, AZ/BNT and AZ/AZ vaccination elicited non-classifiable antibodies that were significantly more potent than their mRNA or Janssen counterparts (Fig. 4b).

To determine the neutralizing breadth of the memory antibodies that developed after AZ/BNT or AZ/AZ vaccination, we analyzed a panel of randomly selected Wuhan-Hu-1 (WT)-neutralizing antibodies from AZ/BNT and AZ/AZ vaccinees (AZ/BNT: n=32, and AZ/AZ: n=34) for neutralizing activity against SARS-CoV-2 pseudoviruses carrying amino acid substitutions specific to the Delta- and Omicron BA.1-RBD.

78% of the AZ/BNT- and 82% of the AZ/AZ-elicited antibodies neutralized SARS-CoV-2 pseudoviruses carrying the Delta RBD-amino acid substitutions, some with IC_50_ values of less than 10 ng/ml (Fig. 4c and Table S4). Omicron BA.1 showed the highest degree of neutralization resistance, nevertheless 8 out of 32 antibodies isolated from AZ/BNT and 14 out of 34 antibodies isolated from AZ/AZ vaccinees neutralized this variant. Some of the most potent Omicron-neutralizing antibodies targeted epitopes that could not be classified in our BLI experiments (non-classified) with IC_50_s below 10 ng/ml (Fig. 4c-d and Table S4). 5 out of 32 AZ/BNT- and 10 out of 34 AZ/AZ- antibodies neutralized both Delta and Omicron, a proportion that was not significantly different compared to antibodies elicited by other vaccine regimens (Fig. 4c and e).

We conclude that the relative distribution of RBD epitopes targeted by neutralizing antibodies expressed by memory B cells that develop after mRNA, Janssen Ad26.COV2.S or ChAdOx1 vaccination regimens differ significantly.

### Structural analysis of antibody-RBD interaction

To understand the interaction between these none-classified antibodies and RBD, we imaged WT Wuhan-Hu-1 SARS-CoV-2 S 6P bound to Fab fragments of a potent and broad AZ/AZ antibody (AZ090) by single-particle cryo-electron microscopy (cryo-EM) (Fig. 5a and Fig. S6). The resolution of the reconstituted cryo-EM electron density map was 3 Å for the whole complex and the spike-AZ090. Structural analyses of the density maps showed that the binding orientation of AZ090 is similar to previously described potent antibodies that were isolated following natural infection (Dejnirattisai et al., 2021; Reincke et al., 2022; Tortorici et al., 2020; Wang et al., 2021a) (Fig. S7). AZ090 and this type of antibodies share the same immunoglobulin heavy and light chain genes (IGHV1-58 and IGKV3-20/IGKJ) (Fig. S7). Unlike Class 1 antibodies, the footprint of AZ90 is located in the ridge region of RBD with more limited overlap with Omicron (BA.1) amino acid substitutions than typical class 1 (C105) and class 2 (C144) antibodies (Fig. 5b). The distinctive binding pattern of AZ090 may also explain the lack of competition in BLI experiments and the neutralizing breadth across different SARS-CoV-2 variants.

**Fig. 5.**
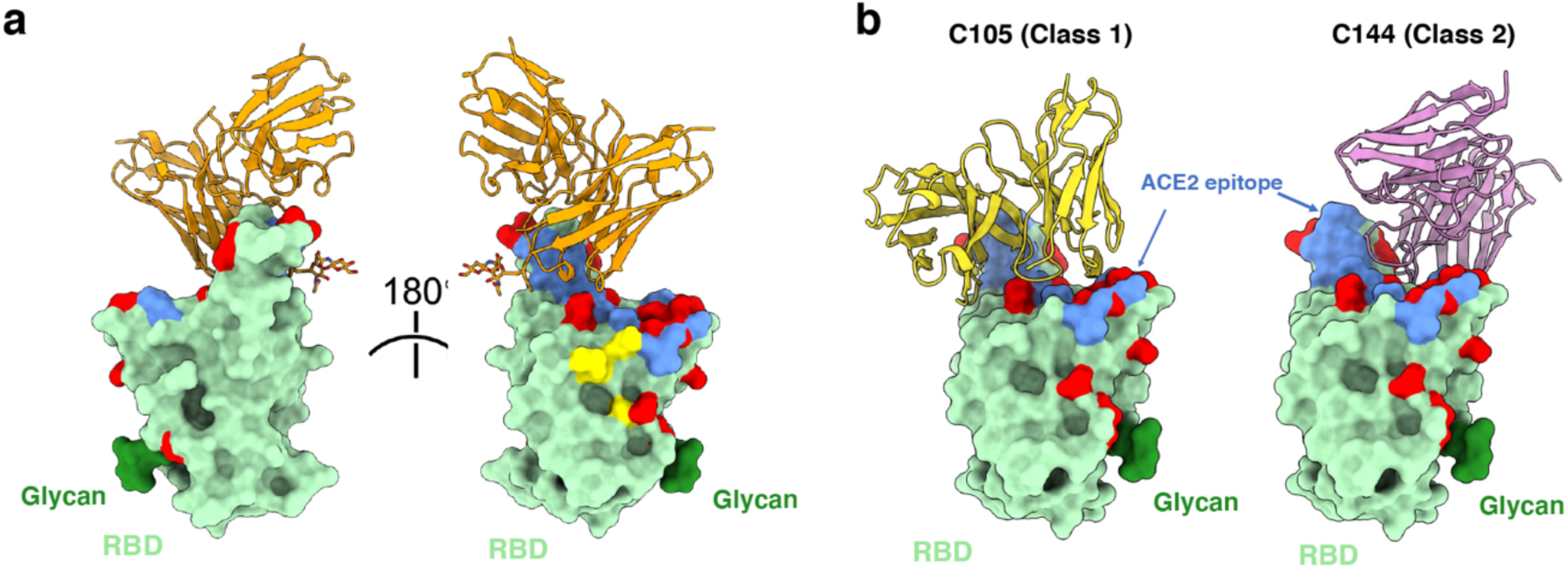
Structural analysis of AZ090 antibody. **a**, RBDs of SARS-CoV-2 were shown by surface and colored green. RBD and AZ090 were shown by cartoon and AZ090 fab was colored in orange, ACE2 epitope was colored blue and N343 glycan was colored green. Omicron (BA.1) mutations were shown red. **b**, as in **a**. C105 (Class 1 antibody, PDB:6XCM) was colored in yellow and C144 (Class 2 antibody, PDB:7K90) was colored in pink.

## Discussion

Neutralizing antibodies are correlates of vaccine efficacy in protection against SARS-CoV-2 infection and severe COVID-19 (Bergwerk et al., 2021; Feng et al., 2021; Khoury et al., 2021; Li et al., 2022). All three US-authorized vaccines have shown substantial protection against SARS-CoV-2 infection, hospitalization, and death (Botton et al., 2022; Self et al., 2021). However, vaccine efficacy wanes over time with prominent loss of protection against infection after the Janssen Ad26.COV2.S vaccine compared to mRNA (Lin et al., 2022). Similarly, vaccination regimens with the globally predominant ChAdOx1vaccine have been less effective in the protection against infection and symptomatic COVID-19 compared to mRNA vaccination (Andrews et al., 2022; Braeye et al., 2022). However, the combination of a ChAdOx1 prime and a 2^nd^ mRNA dose shows similar levels of protection as 2-dose mRNA vaccination (Nordstrom et al., 2021).

Our comparative analysis of plasma and memory B cell antibodies provides a mechanistic explanation for the observed real-world protective efficacy of the several different vaccine regimens. Binding and neutralizing antibody levels elicited by 2-dose mRNA or AZ/BNT vaccination exceeds those elicited by AZ/AZ or single-dose Janssen Ad26.COV2.S vaccination. Omicron BA.1 neutralization was highest after AZ/BNT vaccination suggesting that combination vaccine protocols with extended dosing intervals may induce improved plasma neutralizing responses. In line with our observation, vaccine efficacy has been shown to increase with the interval between the 1^st^ and 2^nd^ vaccine dose (Voysey et al., 2021). Prolonged affinity maturation yielding higher affinity B cells for plasma cell maturation upon the administration of the 2nd vaccine dose may be of importance in this process (Hall et al., 2022).

The relative potency and breadth, i.e. neutralizing activity against Delta and Omicron, of memory B cell antibodies produced by the 4 different vaccine regimens was overall similar. However, they differed in the absolute number of memory cells and the distribution of the RBD epitopes targeted by mRNA, Janssen Ad26.COV2.S and ChAdOx1 vaccination regimens. Differences in dosing intervals between prime and boost immunization, distinct antigenic features of the full-length wild-type SARS-CoV-2 spike protein lacking prefusion-stabilizing mutations in the ChAdOx1 vaccine (Tortorici and Veesler, 2019), and the precise biochemistry of the antigen and its presentation may all contribute to these observations.

Notably, the number of RBD-binding memory cells that develop after 2-dose mRNA vaccination is greater than vaccination regimens that are based on adenoviral vectors. The latter is likely to be particularly important for recall responses and protection from severe diseases upon repeated viral challenge (Amanna et al., 2007; Mesin et al., 2020).

**Fig. S1:**
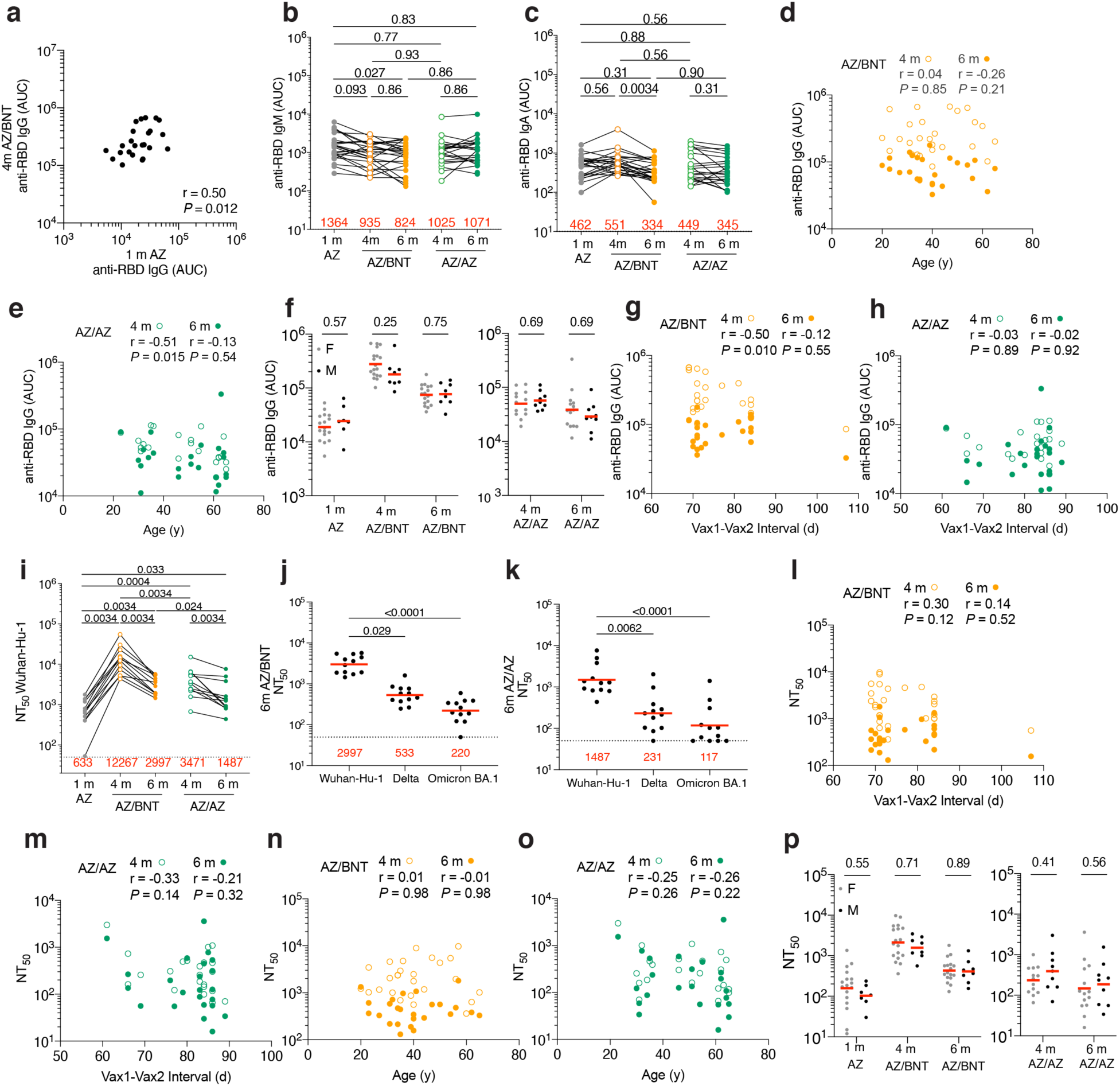
Demographics and Plasma Correlations. **a,** AUC for anti-RBD IgG at 1 m after ChAdOx1(AZ) prime plotted against AUC for anti-RBD IgG at 4 m after initial dose following the ChAdOx1/BNT162b2 AZ/BNT scheme. **b-c**, Area under the curve (AUC) for **b**, plasma IgM and **c**, plasma IgA antibody binding to SARS-CoV-2 Wuhan-Hu-1 RBD 1 months (m) after AZ prime, as well as 4 m and 6 m after initial dose with either BNT162b2 (AZ/BNT; n=26) or ChAdOx1 (AZ/AZ; n=23). Lines connect longitudinal samples. **d-e**, Age (X axis) plotted against area under the curve (AUC) (Y axis) for anti-RBD IgG at 4 m and 6 m after initial dose following **a,** the AZ/BNT scheme, or **b,** the AZ/AZ scheme. **f**, AUC for anti-RBD IgG 1 m after prime, as well as 4 m and 6 m after initial dose for all male (M: n=8) or women (F: n=18) vaccinated following the AZ/BNT scheme (left panel), or AUC for anti-RBD IgG 4 m and 6 m after initial dose for all male (M: n=9) or female (F: n=14) following the AZ/AZ scheme (right panel). **g-h,** Interval between first and second vaccination (X axis) plotted against AUC for anti-RBD IgG (Y axis) at 4 m and 6 m after initial dose following **g,** the AZ/BNT scheme, or **h,** the AZ/AZ scheme. **i-k**, Plasma neutralizing activity against indicated SARS-CoV-2 variants of interest/concern for n=12 randomly selected samples assayed in HT1080Ace2 cl.14 cells. Viruses in i-k contained the R683G furin cleavage site mutation to increase particle infectivity. (**See also in** Fig. 1j**-m**). **l-m**, Interval between first and second vaccination (X axis) plotted against NT_50_ values (Y axis) 4 m and 6 m after initial dose following **l,** the AZ/BNT scheme, or **m,** the AZ/AZ scheme. **n-o**, Age (X axis) plotted against NT_50_ values (Y axis) 4 m and 6 m after initial dose following **n,** the AZ/BNT scheme, or **o,** the AZ/AZ scheme. **p**, NT_50_ values at 1 m after AZ prime, as well as 4 m and 6 m after initial dose for all male (M; n=8) or female (F; n=18) following the AZ/BNT scheme (left panel), or NT_50_ values at 4 m and 6 m after initial dose for all male (M; n=9) or female (F; n=14) following the AZ/AZ scheme (right panel). Red bars represent geometric mean values. r and P values were determined by two-tailed Spearman’s correlation (**d-e, g-h, l-o**). Statistical significance was determined by two-tailed Mann-Whitney test followed by Holm-Šídák test for multiple comparisons (**f, j-k, p**), or by Wilcoxon matched-pairs signed rank test for paired observations followed by Holm-Šídák test for multiple comparisons (**b-c, i**).

**Fig. S2:**
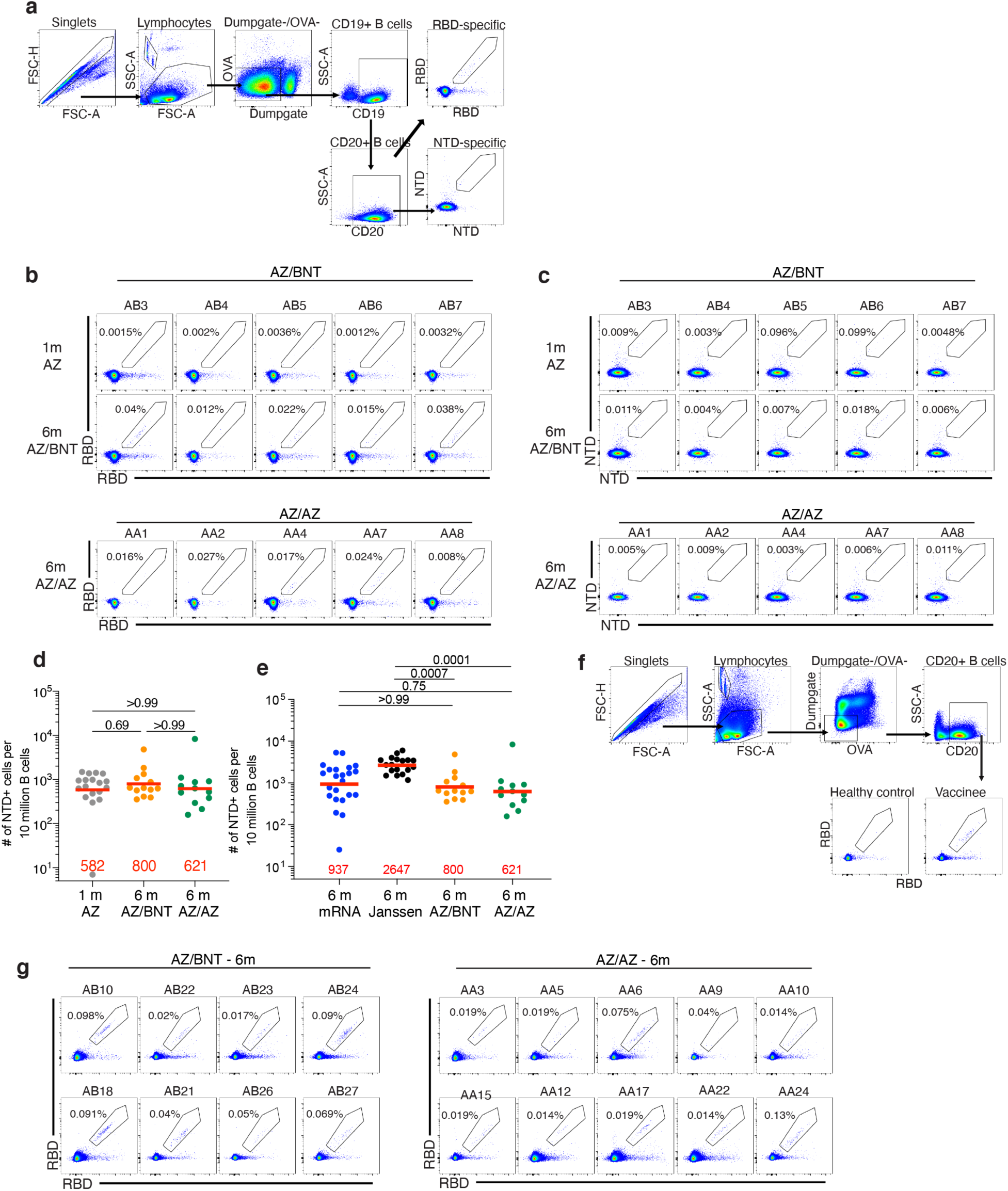
Flow Cytometry. **a**, Gating strategy for phenotyping. Gating was on lymphocytes singlets that were CD19+ or CD20+ and CD3-CD8-CD16-Ova-. Antigen-specific cells were detected based on binding to Wuhan-Hu-1 RBD-PE+ and RBD-AF647+, or to Wuhan-Hu-1 NTD- BrilliantViolet-711+ and NTD- BrilliantViolet-421+. Counting beads were added to each sample and gated based on forward scatter (FSC) and side scatter (SSC) as per manufacturer instructions. **b-c**, Representative flow cytometry plots of **b**, RBD-binding B cells or **c**, NTD-binding B cells in 5 individuals 1 month(m) after AZ prime and 6 months after initial dose. **d-e**, Graph showing the number of NTD-BV711 and NTD-BV421 binding B cells. **f**, Gating strategy for single-cell sorting for CD20+ B cells for RBD-PE and RBD-AF647. **g**, Representative flow cytometry plots showing dual AlexaFluor-647- and PE-Wuhan-Hu-1-RBD binding, single-cell sorted B cells from AZ/BNT and AZ/AZ vaccinees 6m after initial dose.

**Fig. S3:**
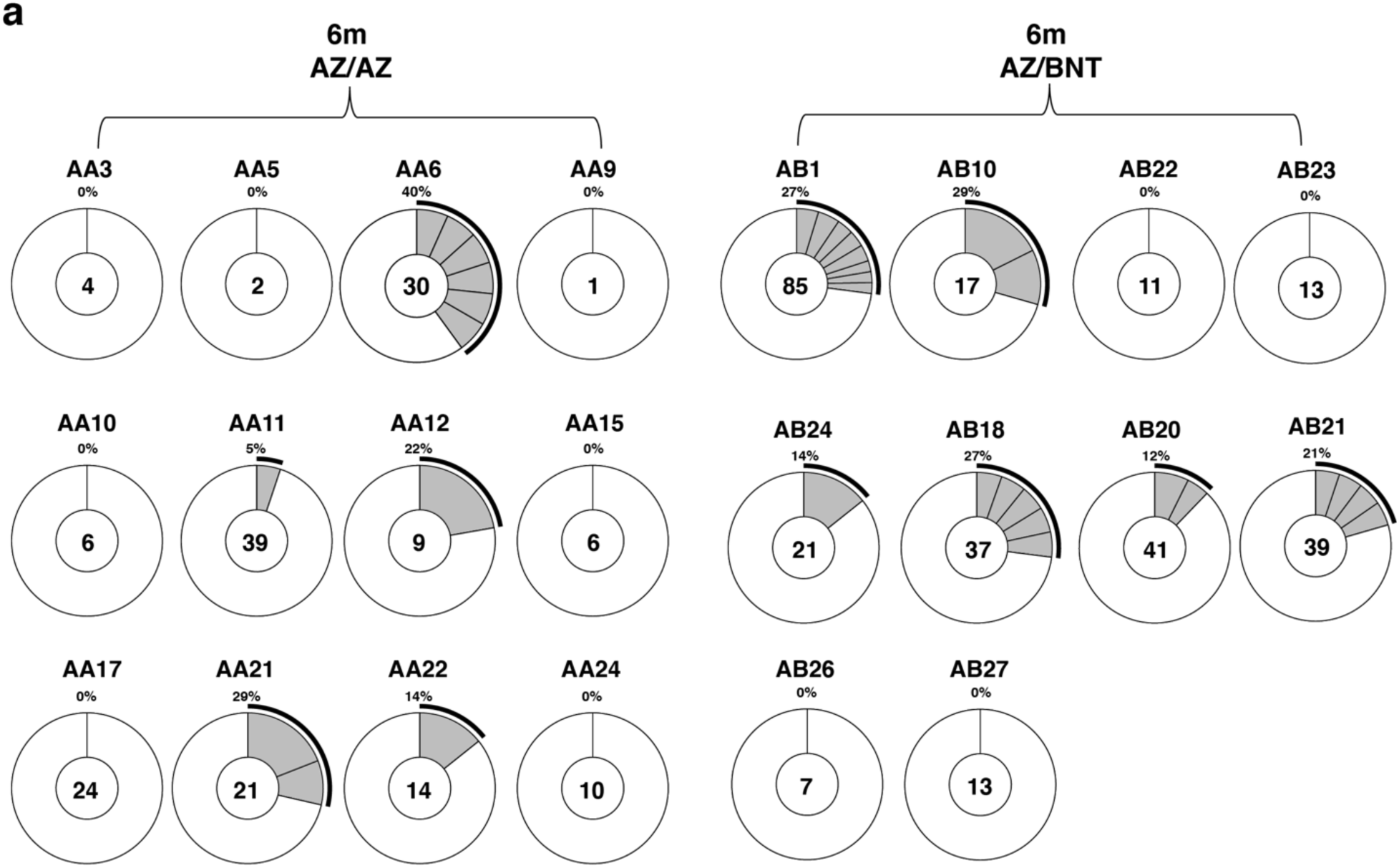
Clonality of anti-SARS-CoV-2 Wuhan-Hu-1 RBD antibody. Pie charts show the distribution of IgG antibody sequences obtained from MBCs from Wuhan- Hu-1 RBD-specific memory B cells of AZ/BNT and AZ/AZ vaccinees 6 m after initial dose. The number inside the circle indicates the number of sequences analyzed for the individual denoted above the circle.

**Fig. S4.**
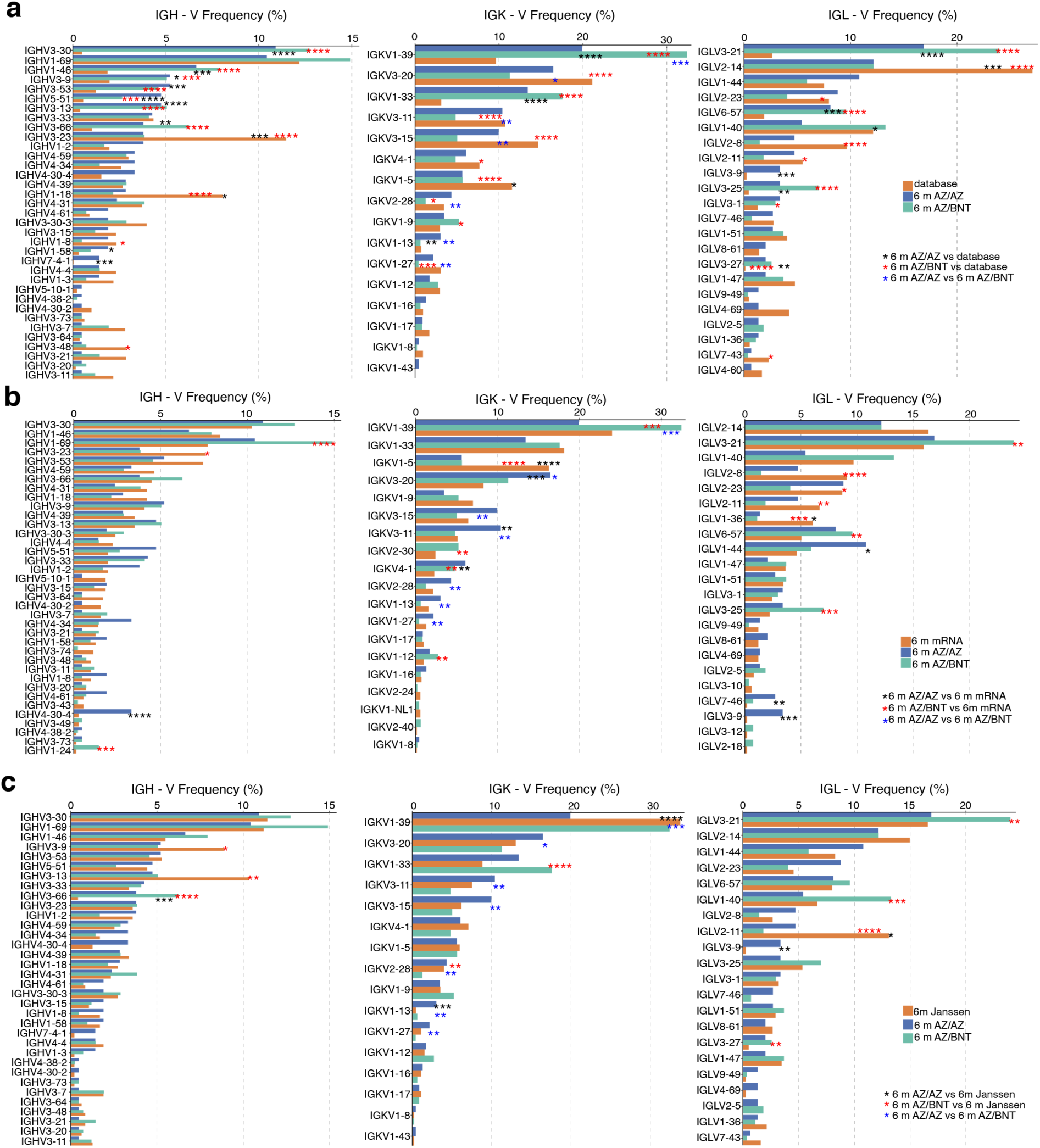
Frequency distribution of human V genes. **a-c** Comparison of the frequency distribution of human V genes for heavy chain and light chains of anti-RBD Wuhan-Hu-1 antibodies from this study and from a database of shared clonotypes of human B cell receptor generated by Cinque Soto et al (Soto et al., 2019). Graph shows relative abundance of human IGHV, IGKV and IGLV genes, with 6 m AZ/AZ antibodies (blue) and AZ/BNT antibodies (green). **a,** Sequence Read Archive accession SRP010970(orange); **b,** antibodies from mRNA vaccinees 6 months(m) after initial dose (orange) (Cho et al., 2021); **c,** antibodies from Janssen Ad26.COV2.S vaccinees 6 m after prime (orange) (Cho et al., 2022). Statistical significance was determined by two-sided binomial test. * = p≤0.05, ** = p≤0.01, *** = p≤0.001, **** = p≤0.0001.

**Fig. S5.**
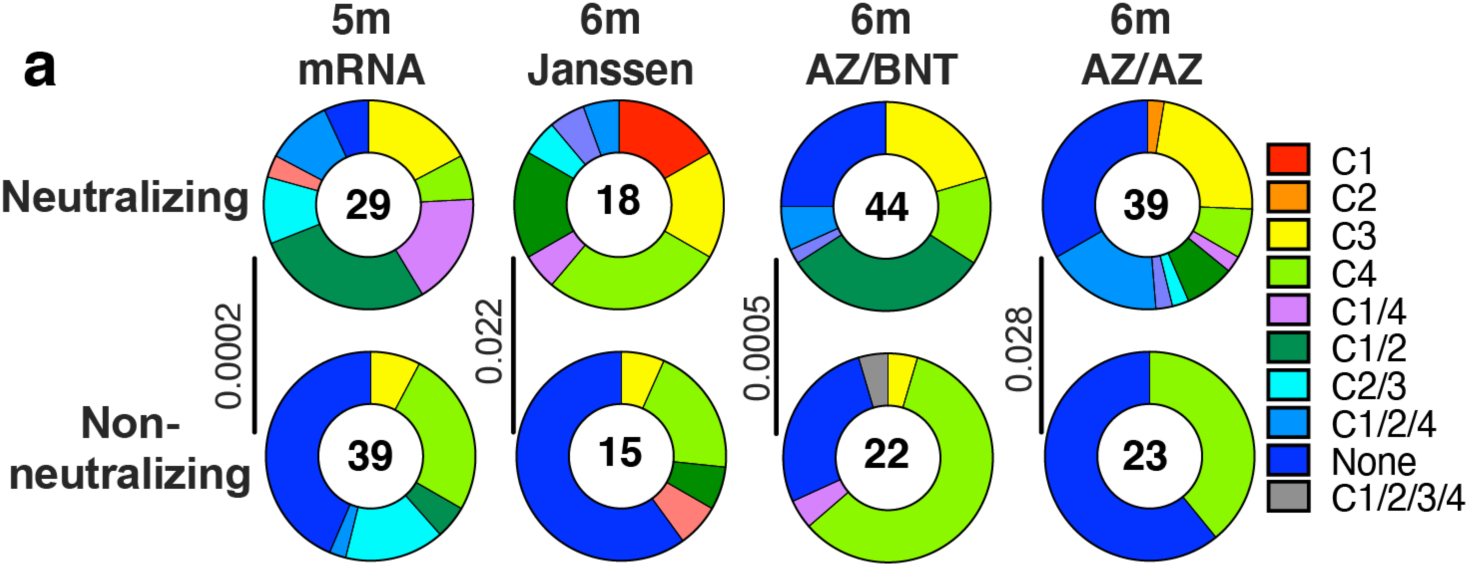
Epitope mapping. **a.** Results of epitope mapping performed by competition BLI. Pie charts show the distribution of the antibody classes among all neutralizing antibodies against Wuhan-Hu-1 and none-neutralizing antibodies obtained from mRNA vaccinees at 6 m after initial dose (n=68) (Cho et al., 2021), Janssen vaccinees at 6 m (n=33) after prime (Cho et al., 2022), to mAbs cloned from AZ/AZ (n=62) or AZ/BNT(n=66) vaccinees 6 m after initial dose. Pie charts were compared using a two-tailed Fisher’s exact test.

**Fig. S6.**
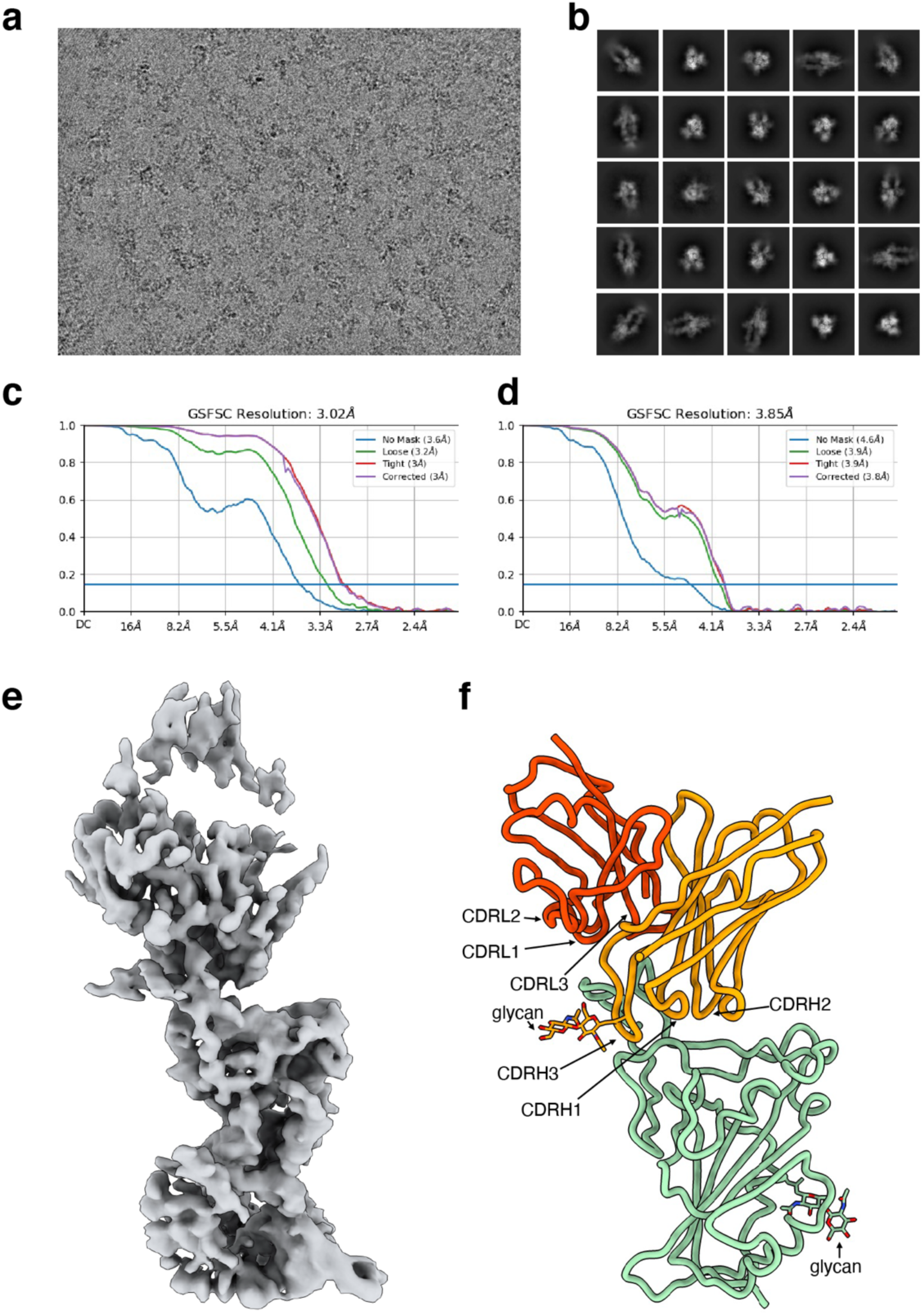
Cryo-EM data processing. **a**, Representative cryo-EM micrograph from whole dataset. **b**, 2D class averages of selected particles for homogeneous refinement. **c**, Gold-standard Fourier shell correlation curves for the whole map of S 6P bound to AZ090 Fabs. **d**, Gold-standard Fourier shell correlation curves for the locally refined reconstruction of the RBD-AZ090 Fab region. **e**, cryo-EM density of RBD-AZ090 Fab region. **f**, Model of Fab fragment bound to RBD of SARS-CoV-2 was shown by cartoon. The glycans were shown by stick. The heavy chain of AZ090 was colored orange and the light chain of AZ090 was colored orange red.

**Fig. S7.**
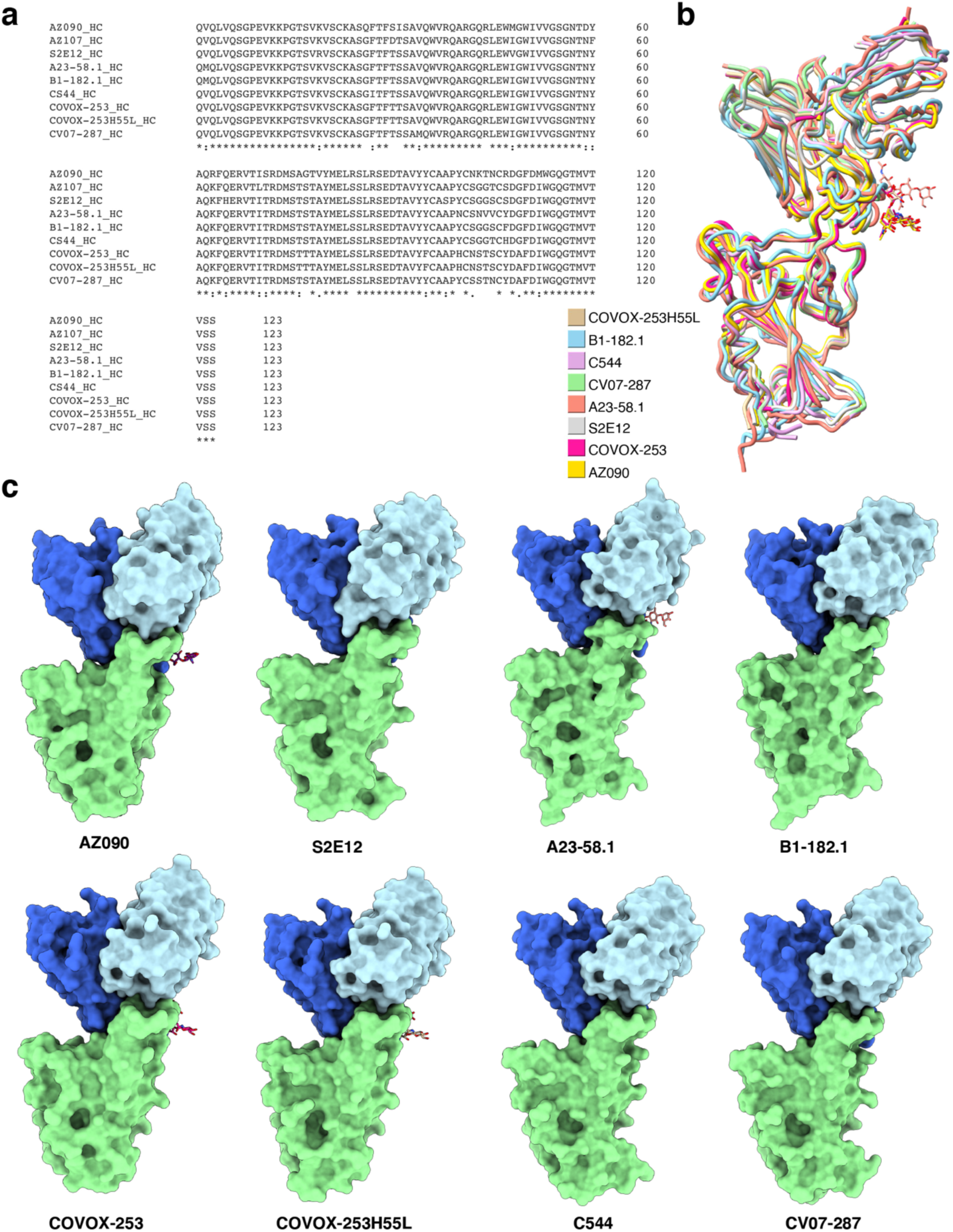
Comparison of AZ090 and previous resolved antibodies. **a**, Multiple sequence alignment of RBDs was processed by Clustal Omega(Sievers et al., 2011). **b**, Structure of AZ090 and previous resolved antibodies encoded by same heavy chains were aligned. Different Fab-RBD structures were colored respectively. **c**, Structures from **b** were shown by cartoon. The RBDs were colored green, the heavy chains were colored royal blue and the light chains were colored light blue. The glycans on the heavy chains were shown by stick.

## Materials and Methods

### Study participants

Health-care workers receiving routine COVID-19 vaccination were enrolled in the EICOV and COVIM prospective observational cohort studies conducted at Charité–Universitätsmedizin Berlin (Berlin, Germany), after written informed consent was obtained. EICOV was approved by the ethics committee of Charité–Universitätsmedizin Berlin (EA4/245/20), and COVIM was approved by the Federal Institute for Vaccines and Biomedicines (Paul Ehrlich Institute) and by the Ethics committee of the state of Berlin (EudraCT-2021–001512–28). Both studies were conducted in accordance with the guidelines of Good Clinical Practice (ICH 1996) and the Declaration of Helsinki. Health-care workers at Charité–Universitätsmedizin Berlin were offered either two doses of BNT162b2 3 weeks apart or an initial dose of ChAdOx1 nCov-19 followed by a heterologous boost with BNT162b2 10–12 weeks later. The vaccine regimen depended on availability and current official recommendations. Health-care workers who received an initial dose of ChAdOx1 nCov-19 were also free to choose a homologous booster with ChAdOx1 nCov-19 10–12 weeks later. (Hillus et al., 2021) For detailed participant characteristics see Table S1 and previous publications (Cho et al., 2021; Cho et al., 2022; Muecksch et al., 2022). Cohort sample analyses were performed under an existing Rockefeller University IRB-approved protocol (DRO-1006).

### Blood samples processing and storage

Blood samples were collected in Heparin and Serum-gel monovette tubes (Greiner bio one). Peripheral Blood Mononuclear Cells (PBMCs) were isolated by gradient centrifugation and stored in liquid nitrogen in the presence of Fetal Calf Serum (FCS) and Dimethylsulfoxide (DMSO). Heparinized plasma and serum samples were fractioned by centrifugation, aliquoted and stored at -80°C until analysis. Prior to experiments, aliquots of plasma samples were heat-inactivated (56°C for 30 minutes) and then stored at 4°C.

### ELISAs

Enzyme-Linked Immunosorbent Assays (ELISAs) (Amanat et al., 2020; Grifoni et al., 2020) to evaluate antibodies binding to SARS-CoV-2 RBD were performed by coating of high-binding 96-half-well plates (Corning 3690) with 50 μl per well of a 1μg/ml protein solution in Phosphate-buffered Saline (PBS) overnight at 4°C. Plates were washed 6 times with washing buffer (1× PBS with 0.05% Tween-20 (Sigma-Aldrich)) and incubated with 170 μl per well blocking buffer (1× PBS with 2% BSA and 0.05% Tween-20 (Sigma)) for 1 hour at room temperature.

Immediately after blocking, monoclonal antibodies or plasma samples were added in PBS and incubated for 1 hour at room temperature. Plasma samples were assayed at a 1:66 starting dilution and 10 additional threefold serial dilutions. Monoclonal antibodies were tested at 10 μg/ml starting concentration and 10 additional fourfold serial dilutions. Plates were washed 6 times with washing buffer and then incubated with anti-human IgG, IgM or IgA secondary antibody conjugated to horseradish peroxidase (HRP) (Jackson ImmunoResearch 109-036-088, 109-035-129, and Sigma A0295) in blocking buffer at a 1:5,000 dilution (IgM and IgG) or 1:3,000 dilution (IgA). Plates were developed by addition of the HRP substrate, 3,3’,5,5’- Tetramethylbenzidine (TMB) (ThermoFisher) for 10 minutes (plasma samples) or 4 minutes (monoclonal antibodies). The developing reaction was stopped by adding 50 μl of 1 M H_2_SO_4_ and absorbance was measured at 450 nm with an ELISA microplate reader (FluoStar Omega, BMG Labtech) with Omega and Omega MARS software for analysis. For plasma samples, a positive control (plasma from participant COV72, diluted 66.6-fold and ten additional threefold serial dilutions in PBS) was added to every assay plate for normalization. The average of its signal was used for normalization of all the other values on the same plate with Excel software before calculating the area under the curve using Prism V9.1 (GraphPad). Negative controls of pre-pandemic plasma samples from healthy donors were used for validation (for more details, please see (Robbiani et al., 2020)). For monoclonal antibodies, the ELISA half-maximal concentration (EC_50_) was determined using four-parameter nonlinear regression (GraphPad Prism V9.1). EC_50_s above 1000 ng/mL were considered non-binders.

### Proteins

The mammalian expression vector encoding the Receptor Binding-Domain (RBD) of SARS-CoV-2 (GenBank MN985325.1; Spike (S) protein residues 319-539) was previously described (Barnes et al., 2020). Mammalian expression vector encoding the SARS-CoV-2 Wuhan-Hu-1 NTD (GenBank MN985325.1; S protein residues 14-307) was previously described (Wang et al., 2022a).

### SARS-CoV-2 pseudotyped reporter virus

A panel of plasmids expressing RBD-mutant SARS-CoV-2 spike proteins in the context of pSARS-CoV-2-S _Δ19_ has been described (Cho et al., 2021; Muecksch et al., 2021; Wang et al., 2021d; Weisblum et al., 2020). Variant pseudoviruses resembling SARS-CoV-2 variants Delta (B.1.617.2) and Omicron BA.1 (B.1.1.529) have been described before (Cho et al., 2021; Schmidt et al., 2021a; Wang et al., 2021c) and were generated by introduction of substitutions using synthetic gene fragments (IDT) or overlap extension PCR mediated mutagenesis and Gibson assembly. Specifically, the variant-specific deletions and substitutions introduced were: Delta: T19R, Δ156-158, L452R, T478K, D614G, P681R, D950N Delta-RBD: L452R, T478K Omicron BA.1: A67V, Δ69-70, T95I, G142D, Δ143-145, Δ211, L212I, ins214EPE, G339D, S371L, S373P, S375F, K417N, N440K, G446S, S477N, T478K, E484A, Q493K, G496S, Q498R, N501Y, Y505H, T547K, D614G, H655Y, H679K, P681H, N764K, D796Y, N856K, Q954H, N969H, N969K, L981F Deletions/substitutions corresponding to variants of concern listed above, were incorporated into a spike protein that also includes the R683G substitution, which disrupts the furin cleavage site and increases particle infectivity. Neutralizing activity against mutant pseudoviruses were compared to a wildtype (WT) SARS-CoV-2 spike sequence (NC_045512), carrying R683G where appropriate.

SARS-CoV-2 pseudotyped particles were generated as previously described (Robbiani et al., 2020; Schmidt et al., 2020). Briefly, 293T (CRL-11268) cells were obtained from ATCC, and the cells were transfected with pNL4-3 ΔEnv-nanoluc and pSARS-CoV-2-S_Δ19_. Particles were harvested 48 hours post-transfection, filtered and stored at -80°C.

### Pseudotyped virus neutralization assay

Four- to five-fold serially diluted pre-pandemic negative control plasma from healthy donors, plasma from individuals who received Ad26.COV2.S vaccines, or monoclonal antibodies were incubated with SARS-CoV-2 pseudotyped virus for 1 hour at 37 °C. The mixture was subsequently incubated with 293T_Ace2_ cells (Robbiani et al., 2020) (for all WT neutralization assays) or HT1080Ace2 cl14 (for all mutant panels and variant neutralization assays) cells (Wang et al., 2021d) for 48 hours after which cells were washed with PBS and lysed with Luciferase Cell Culture Lysis 5× reagent (Promega). Nanoluc Luciferase activity in lysates was measured using the Nano-Glo Luciferase Assay System (Promega) with the Glomax Navigator (Promega) or ClarioStar Microplate Multimode Reader (BMG). The relative luminescence units were normalized to those derived from cells infected with SARS-CoV-2 pseudotyped virus in the absence of plasma or monoclonal antibodies. The half-maximal neutralization titers for plasma (NT_50_) or half-maximal and 90% inhibitory concentrations for monoclonal antibodies (IC_50_ and IC_90_) were determined using four-parameter nonlinear regression (least squares regression method without weighting; constraints: top=1, bottom=0) (GraphPad Prism).

### Biotinylation of viral protein for use in flow cytometry

Purified and Avi-tagged SARS-CoV-2 Wuhan-Hu-1 RBD and NTD were biotinylated using the Biotin-Protein Ligase-BIRA kit according to manufacturer’s instructions (Avidity) as described before (Robbiani et al., 2020). Ovalbumin (Sigma, A5503-1G) was biotinylated using the EZ-Link Sulfo-NHS-LC-Biotinylation kit according to the manufacturer’s instructions (Thermo Scientific). Biotinylated ovalbumin was conjugated to streptavidin-BB515 (BD, 564453). RBD was conjugated to streptavidin-PE (BD Biosciences, 554061) and streptavidin-AF647 (Biolegend, 405237) (Robbiani et al., 2020). NTD was conjugated to streptavidin-BV421 (Biolegend, 405225) and streptavidin-BV711 (BD Biosciences, 563262).

### Flow cytometry and single cell sorting

Single-cell sorting by flow cytometry was described previously (Robbiani et al., 2020). Briefly, peripheral blood mononuclear cells were enriched for B cells by negative selection using a pan- B-cell isolation kit according to the manufacturer’s instructions (Miltenyi Biotec, 130-101-638). The enriched B cells were incubated in Flourescence-Activated Cell-sorting (FACS) buffer (1× PBS, 2% FCS, 1 mM ethylenediaminetetraacetic acid (EDTA)) with the following anti-human antibodies (all at 1:200 dilution): anti-CD20-PECy7 (BD Biosciences, 335793), anti-CD3-APC- eFluro780 (Invitrogen, 47-0037-41), anti-CD8-APC-eFluor780 (Invitrogen, 47-0086-42), anti- CD16-APC-eFluor780 (Invitrogen, 47-0168-41), anti-CD14-APC-eFluor780 (Invitrogen, 47- 0149-42), as well as Zombie NIR (BioLegend, 423105) and fluorophore-labeled Wuhan-Hu-1 RBD, NTD, and ovalbumin (Ova) for 30 min on ice. AccuCheck Counting Beads (Life Technologies, PCB100) were added to each sample according to manufacturer’s instructions.

Single CD3-CD8-CD14-CD16−CD20+Ova− B cells that were RBD-PE+RBD-AF647+ were sorted into individual wells of 96-well plates containing 4 μl of lysis buffer (0.5× PBS, 10 mM Dithiothreitol (DTT), 3,000 units/ml RNasin Ribonuclease Inhibitors (Promega, N2615)) per well using a FACS Aria III and FACSDiva software (Becton Dickinson) for acquisition and FlowJo for analysis. The sorted cells were frozen on dry ice, and then stored at −80 °C or immediately used for subsequent RNA reverse transcription.

### Antibody sequencing, cloning and expression

Antibodies were identified and sequenced as described previously (Robbiani et al., 2020; Wang et al., 2021b). In brief, RNA from single cells was reverse-transcribed (SuperScript III Reverse Transcriptase, Invitrogen, 18080-044) and the cDNA was stored at −20 °C or used for subsequent amplification of the variable IGH, IGL and IGK genes by nested PCR and Sanger sequencing. Sequence analysis was performed using MacVector. Amplicons from the first PCR reaction were used as templates for sequence- and ligation-independent cloning into antibody expression vectors. Recombinant monoclonal antibodies were produced and purified as previously described (Robbiani et al., 2020).

### Biolayer interferometry

Biolayer interferometry assays were performed as previously described (Robbiani et al., 2020). Briefly, we used the Octet Red instrument (ForteBio) at 30°C with shaking at 1,000 r.p.m.

Epitope binding assays were performed with protein A biosensor (ForteBio 18-5010), following the manufacturer’s protocol “classical sandwich assay” as follows: (1) Sensor check: sensors immersed 30 sec in buffer alone (buffer ForteBio 18-1105), (2) Capture 1st Ab: sensors immersed 10 min with Ab1 at 10 µg/mL, (3) Baseline: sensors immersed 30 sec in buffer alone, (4) Blocking: sensors immersed 5 min with IgG isotype control at 10 µg/mL. (5) Baseline: sensors immersed 30 sec in buffer alone, (6) Antigen association: sensors immersed 5 min with RBD at 10 µg/mL. (7) Baseline: sensors immersed 30 sec in buffer alone. (8) Association Ab2: sensors immersed 5 min with Ab2 at 10 µg/mL. Affinity measurement of anti-SARS-CoV-2 IgGs binding were corrected by subtracting the signal obtained from traces performed with IgGs in the absence of RBD. The kinetic analysis using protein A biosensor (ForteBio 18-5010) was performed as follows: (1) baseline: 60sec immersion in buffer. (2) loading: 200sec immersion in a solution with IgGs 10 μg/ml. (3) baseline: 200sec immersion in buffer. (4) Association: 300sec immersion in solution with RBD at 20, 10, or 5 μg/ml (5) dissociation: 600sec immersion in buffer. Curve fitting was performed using a fast 1:1 binding model and the Data analysis software (ForteBio). Mean *K*_D_ values were determined by averaging all binding curves that matched the theoretical fit with an R^2^ value ≥ 0.8.Curve fitting was performed using the Fortebio Octet Data analysis software (ForteBio).

### Recombinant protein expression

Stabilized SARS-CoV-2 6P ectodomain and Fabs were expressed and purified as previously described(Wang et al., 2022b). Briefly, constructs encoding the stabilized spike of SARS-CoV-2 ectodomain(Hsieh et al., 2020) were used to transiently transfect Expi293F cells (Gibco).

Supernatants were harvested after four days, and S 6P proteins were purified by nickel affinity following with size-exclusion chromatography. Peak fractions from size-exclusion chromatography were identified by native gel analysis for spike trimer fractions.

### Cryo-EM sample preparation

Purified Fabs were mixed with S 6P protein at a 1.1:1 M ratio of Fab-to-protomer for 30 min at room temperature. Fab-S complexes were deposited on a freshly glow-discharged 400 mesh, 1.2/1.3 Quantifoil grid (Electron Microscopy Sciences). Immediately prior to deposition of 3 mL of complex onto grid, fluorinated octyl-maltoside (Anatrace) was added to the sample to a final concentration of 0.02% w/v. Samples were vitrified in 100% liquid ethane using a Mark IV Vitrobot (Thermo Fisher) after blotting at 22 C° and 100% humidity for 3s with filter paper.

### Cryo-EM data collection and processing

Single-particle cryo-EM data were collected on a Titan Krios transmission electron microscope (Thermo Fisher) equipped with a Gatan K3 direct detector, operating at 300 kV and controlled using SerialEM automated data collection software (Mastronarde, 2005). A total dose of 56.56 e/Å^2^ was accumulated on each movie with a pixel size of 0.515 and a defocus range of -0.8 and 2.0 µm. Movie frame alignment, CTF estimation, particle-picking and extraction were carried out using cryoSPARC v3.3.1(Punjani et al., 2017). Reference-free particle picking and extraction were performed on dose-weighted micrographs. A subset of 4x-downsampled particles were used to conduct several rounds of reference-free 2D classification, then the selected Fab-S particles were extracted, 2x-downsampled, yielding a pixel size of 1.03 Å. The particles were used to generate *ab initio* models, which were then used for heterogeneous refinement of the entire dataset in cryoSPARC. Particles belonging to classes that resembled Fab-S structures were homogeneous refined following with non-uniform refinement until imported into Relion 3.1.3 for CTF refinement. The particles were then imported into cryoSPARC for heterogenerous refinement. Particles belonging to classes with better Fab density were selected and subjected to another round of homogeneous refinement following with non-uniform refinement. To improve the density of the RBD/AZ090interface, several rounds of local refinement were then performed using different soft masks. Reported resolutions are based on the gold-standard Fourier shell correlation of 0.143 criterion (Bell et al., 2016).

### Cryo-EM structure modeling and analysis

UCSF Chimera(Pettersen et al., 2004) and Coot(Emsley et al., 2010) were used to fit atomic models into the locally refined cryoEM map. Models were refined and validated by Phenix(Liebschner et al., 2019). Figures were generated using UCSF ChimeraX(Goddard et al., 2018).

### Computational analyses of antibody sequences

Antibody sequences were trimmed based on quality and annotated using Igblastn v.1.14. with IMGT domain delineation system. Annotation was performed systematically using Change-O toolkit v.0.4.540 (Gupta et al., 2015). Clonality of heavy and light chain was determined using DefineClones.py implemented by Change-O v0.4.5 (Gupta et al., 2015). The script calculates the Hamming distance between each sequence in the data set and its nearest neighbor. Distances are subsequently normalized and to account for differences in junction sequence length, and clonality is determined based on a cut-off threshold of 0.15. Heavy and light chains derived from the same cell were subsequently paired, and clonotypes were assigned based on their V and J genes using in-house R and Perl scripts. All scripts and the data used to process antibody sequences are publicly available on GitHub (https://github.com/stratust/igpipeline/tree/igpipeline2_timepoint_v2).

The frequency distributions of human V genes in anti-SARS-CoV-2 antibodies from this study was compared to 131,284,220 IgH and IgL sequences generated by (Soto et al., 2019) and downloaded from cAb-Rep (Guo et al., 2019), a database of human shared BCR clonotypes available at https://cab-rep.c2b2.columbia.edu/. Based on the 150 distinct V genes that make up the 1099 analyzed sequences from Ig repertoire of the 6 participants present in this study, we selected the IgH and IgL sequences from the database that are partially coded by the same V genes and counted them according to the constant region. The frequencies shown in Fig. S4 are relative to the source and isotype analyzed. We used the two-sided binomial test to check whether the number of sequences belonging to a specific *IGHV* or *IGLV* gene in the repertoire is different according to the frequency of the same IgV gene in the database. Adjusted p-values were calculated using the false discovery rate (FDR) correction. Significant differences are denoted with stars.

Nucleotide somatic hypermutation and Complementarity-Determining Region 3 (CDR3) length were determined using in-house R and Perl scripts. For somatic hypermutations (SHM), *IGHV* and *IGLV* nucleotide sequences were aligned against their closest germlines using Igblastn and the number of differences were considered nucleotide mutations. The average number of mutations for V genes was calculated by dividing the sum of all nucleotide mutations across all participants by the number of sequences used for the analysis.

## Data presentation

Figures arranged in Adobe Illustrator 2022.

## Data availability statement

Data are provided in Tables S1-5. The raw sequencing data and computer scripts associated with Fig. 2 have been deposited at Github (https://github.com/stratust/igpipeline/tree/igpipeline2_timepoint_v2). This study also uses data from (DeWitt et al., 2016), cAb-Rep (https://cab-rep.c2b2.columbia.edu/)(Guoet al., 2019), Sequence Read Archive (accession SRP010970), and from (Soto et al., 2019).

## Code availability statement

Computer code to process the antibody sequences is available at GitHub (https://github.com/stratust/igpipeline/tree/igpipeline2_timepoint_v2).

## Acknowledgements

We thank all study participants who devoted time to our research, The Rockefeller University Hospital nursing staff and Clinical Research Support Office. We thank Dr. Linas Urnavicius for the help on cryo-EM data processing. Cryo-EM data for this work was collected at the Evelyn Gruss Lipper cryo-EM resource center (the Rockefeller University) with support from Mark Ebrahim, Johanna Sotiris, and Honkit Ng. We thank all members of the M.C.N. laboratory for helpful discussions, Maša Jankovic for laboratory support and Kristie Gordon for technical assistance with cell-sorting experiments. We thank Stefanie Jentzsch and all members of the COVIMMUNIZE/COVIM Study Group for sample acquisition and processing. This work was supported by NIH grant P01-AI138398-S1 (M.C.N.) and 2U19AI111825 (M.C.N.). R37-AI64003 to P.D.B.; R01AI78788 to T.H.; P01AI165075 to M.C.N., P.D.B and T.H.; COVIM: NaFoUniMedCovid19 (FKZ: 01KX2021) to LES, the Federal Institute for Drugs and Medical Devices (V-2021.3 / 1503_68403 / 2021-2022) to FK and LES, the Deutsche Forschungsgemeinschaft (DFG) SFB-TR84 to LES and a donation from Zalando SE to Charité - Universitätsmedizin Berlin.

F.M. was supported by the Bulgari Women and Science Fellowship for COVID-19 Research. C.G. was supported by the Robert S. Wennett Post-Doctoral Fellowship, in part by the National Center for Advancing Translational Sciences (National Institutes of Health Clinical and Translational Science Award program, grant UL1 TR001866), and by the Shapiro-Silverberg Fund for the Advancement of Translational Research. P.D.B. and M.C.N. are Howard Hughes Medical Institute Investigators. This article is subject to HHMI’s Open Access to Publications policy. HHMI lab heads have previously granted a nonexclusive CC BY 4.0 license to the public and a sublicensable license to HHMI in their research articles. Pursuant to those licenses, the author-accepted manuscript of this article can be made freely available under a CC BY 4.0 license immediately upon publication.

## Author information

Z.W., F.Muecksch, F.Muenn and A.C. contributed equally to this work.

## Author Contributions

Z.W., F.M., A.C., T.H., P.D.B., M.C.N. and C.G. conceived, designed, and analyzed the experiments. F.K., L.S., M. C. and C.G. designed clinical protocols. F.M. Z.W., A.C., S.H., R.R., T.B.T, J.D., E.B., C.G.-C., K.Y., and C.G. carried out experiments. B.J. and A.G., produced antibodies. A.G. produced SARS-CoV-2 proteins. F.Muenn, P.T.-L., D.H., F.K., L.S., M.T., K.G.M., I.S., C.G. and M.C. recruited participants, executed clinical protocols, and processed samples. T.Y.O. and V.R. performed bioinformatic analysis. Z.W., M.C.N. and C.G. wrote the manuscript with input from all co-authors.

## Corresponding authors

Correspondence should be addressed to Michel C. Nussenzweig or Christian Gaebler

